# Evolution of the standard genetic code

**DOI:** 10.1101/2020.02.20.958546

**Authors:** Michael Yarus

**Affiliations:** Department of Molecular, Cellular and Developmental Biology, University of Colorado Boulder, Boulder, CO 80309-0347

**Keywords:** coding table, codon, triplet, evolution, distribution fitness

## Abstract

A near-universal Standard Genetic Code (SGC) implies a single origin for Earthly life. To study this unique event, I compute paths to the SGC, comparing different plausible histories. Notably, SGC-like coding emerges from traditional evolutionary mechanisms, and a superior path can be identified.

To objectively measure evolution, progress values from 0 (random coding) to 1 (SGC-like) are defined: these measure fractions of random-code-to-SGC distance. Progress types are **spacing**/**distance**/**d**elta **P**olar **R**equirement, detecting space between identical assignments /mutational distance to the SGC/chemical order, respectively. A coding system was based on known RNAs performing aminoacyl-RNA synthetase reactions. Acceptor RNAs exhibit SGC-like wobble; alternatively, non-wobbling triplets uniquely encode 20 amino acids/start/stop. Triplets acquire 22 functions by stereochemistry, selection, coevolution, or randomly. Assignments also propagate to an assigned triplet’s neighborhood via single mutations, but can also decay.

Futile evolutionary paths are plentiful due to the vast code universe. Thus SGC evolution is critically sensitive to disorder from random assignments. Evolution also inevitably slows near coding completion. Coding likely avoided these difficulties, and two suitable paths are compared. In *late wobble*, a majority of non-wobble assignments are made before wobble is adopted. In *continuous wobble*, a uniquely advantageous early intermediate supplies the gateway to an ordered SGC. Revised coding evolution (limited randomness, late wobble, concentration on amino acid encoding, chemically conservative coevolution with a chemically-ordered elite) produces varied full codes with excellent joint progress values. A population of only 600 independent coding tables includes SGC-like members; a Bayesian path toward more accurate SGC evolution is available.

## Introduction

### The object of the investigation

Fig. 1A initiates analysis by depicting its goal. The figure contains the SGC, connecting codon triplets and standard abbreviations for encoded functions, like the 20 standard amino acids. (Woese 1965) discovered that the chromatographic mobility of amino acids in organic heterocycle/water mixed solvents could be used to classify the amino acids in a way relevant to the genetic code. In particular, the dependence of chromatographic mobility on the mole fraction water in the mixed solvent, called the ‘polar requirement’, has been attached in parentheses to the amino acid abbreviations in Fig. 1A. Here polar requirements are not Woese’s original chromatographic values, but these quantities corrected (Mathew and Luthey-Schulten 2008) by molecular dynamics distribution studies, which can circumvent chromatographic artifacts, such as amino acids with affinity for a paper chromatographic support.

**Figure 1A.**
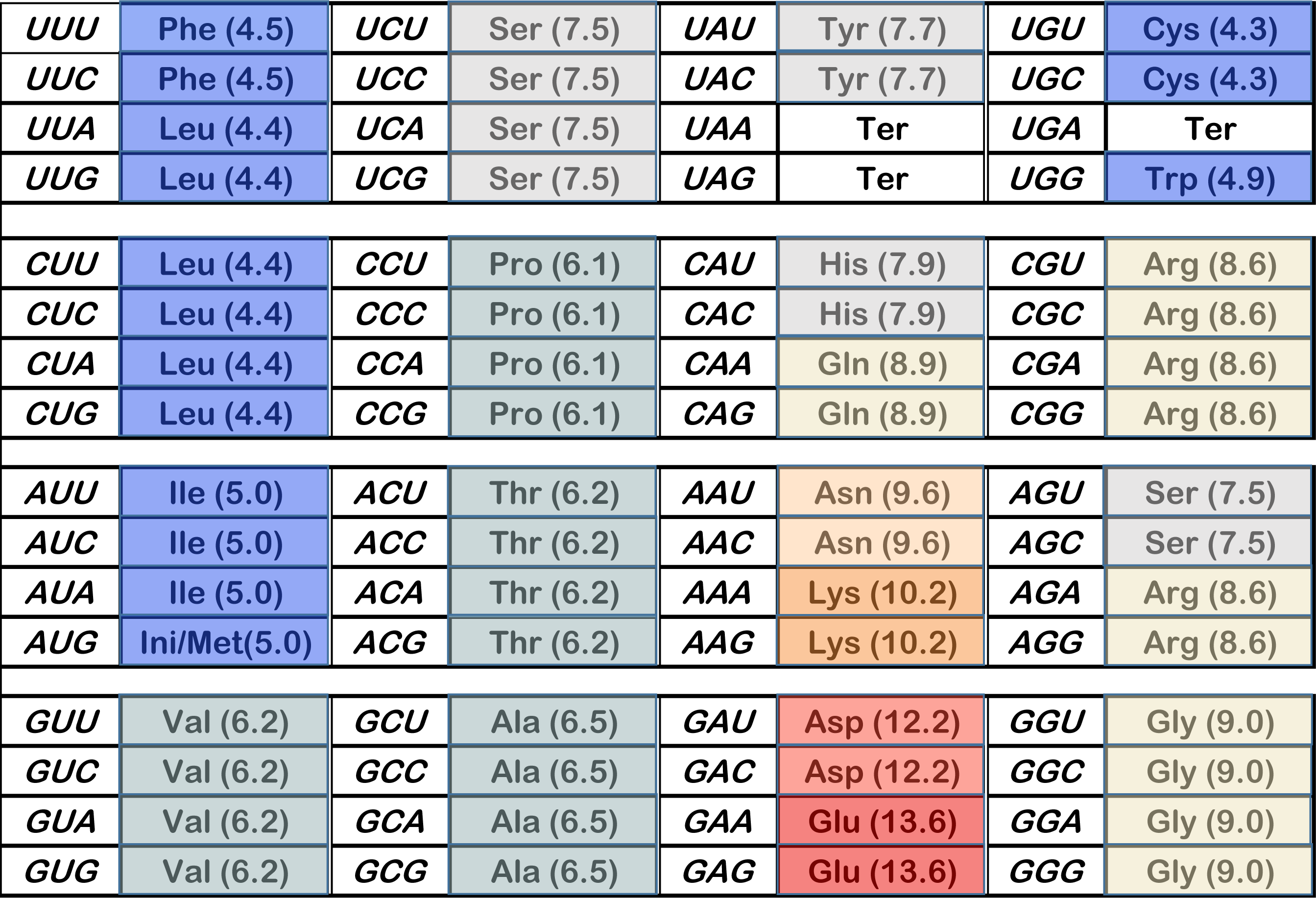
**bottom**. The standard genetic code (SGC), with parenthetical polar requirements (Mathew and Luthey-Schulten 2008). The SGC has progress values: Spacing = 1.0, Distance = 1.0; dPR = 1.0

**Figure 1A.**
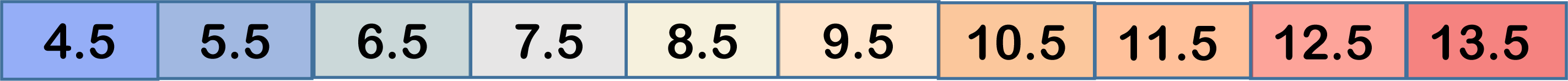
**top**. Color coding for polar requirement in Fig. 1. Each number indicates the midpoint PR for that color. So: the 10.5 box spans 10.01 to 11.0.

Woese pointed out (Woese et al. 1966) that the genetic code assigned similar codons to amino acids with similar polar requirements. In Fig. 1A each triplet has been colored, with hydrophobic polar requirements blue, intermediate ones gray to beige, and very polar side chains red. The SGC is exceedingly highly ordered with respect to the polar requirement, with large coherent domains for hydrophobic, intermediate, and polar amino acids. The single isolated chemical domain is also the smallest; at the upper right, containing the unusual amino acids Cys and Trp. Chemical order spans the coding table, with division into a few coherent regions especially striking. This coherence makes it obvious why the code’s development can be accurately directed by maximizing similarity in polar requirements as a guide (Freeland and Hurst 1998b). This has been attributed to similar roles for chemically similar amino acids within proteins (but see below).

To illustrate extent of SGC order by contrast, Fig. 1B is a coding table that has none. Triplets were assigned using randomized numbers, then the table was colored using the polar requirement scheme of Fig. 1A. The distinctive, pervasive chemical order of the SGC is strikingly evident in the dissimilarity of Fig. 1A and B.

**Figure 1B.**
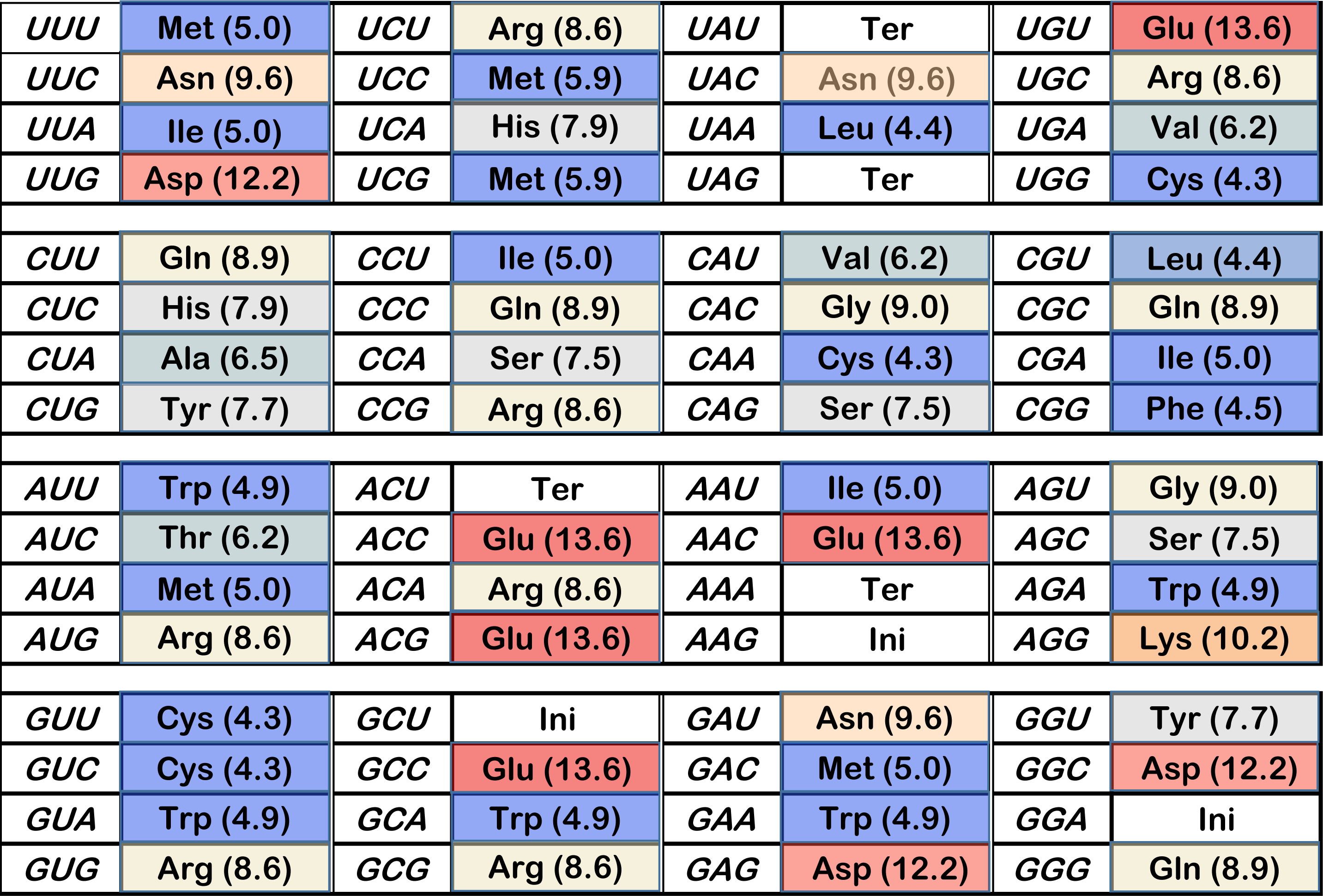
A randomized genetic coding table; each triplet assigned to one of 22 functions by randomized number. Colors visually represent polar requirement, as in Fig. 1A. The example shown is representative of randomized coding tables: its progress values: Spacing = 0.009, Distance = -0.017, dPR = -0.195.

### A model for calculations

To investigate SGC appearance, we desire the fewest, least specific assumptions, in order to maximally respect limited knowledge of the early code. These are: there was an era in which 22 meanings (20 amino acids and start and stop signals) became assigned to 64 possible triplets. This era begins with the first triplet assignment, and ends with a fully assigned coding table that resembles the SGC (**S**tandard **G**enetic **C**ode). Meaningful average rates of coding assignment, which includes both enabling mutation and ensuing events that fix a new meaning, are assumed to exist (Methods, Fig. 12).

**Figure 2A.**
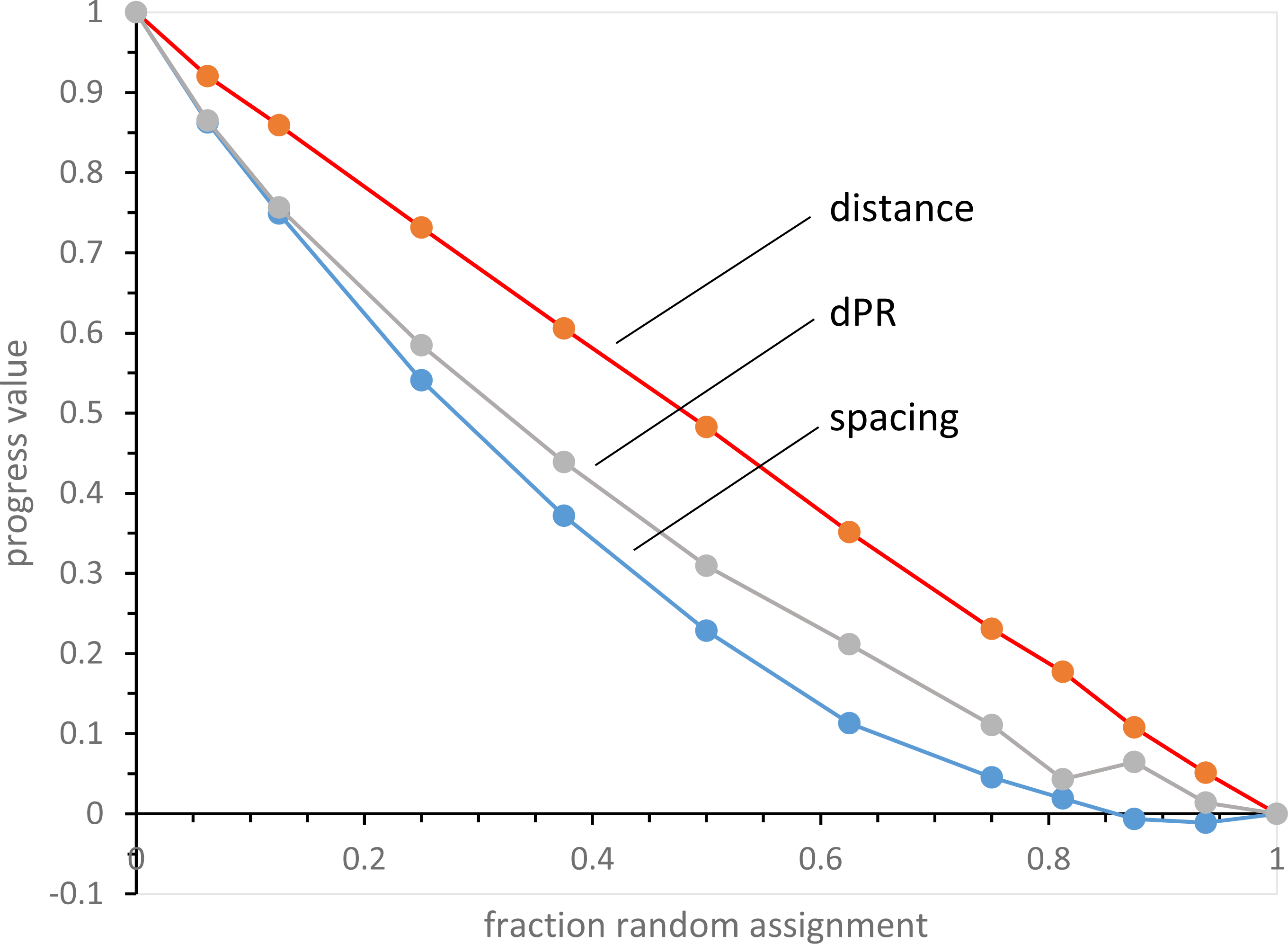
Progress values in randomly assembled coding tables. Computed mean spacing, distance and dPR progress values for 250 full coding tables at each point, assembled with the abscissa’s fraction of varied random assignments, complemented with SGC-like assignments for all other triplets.

**Figure 2B.**
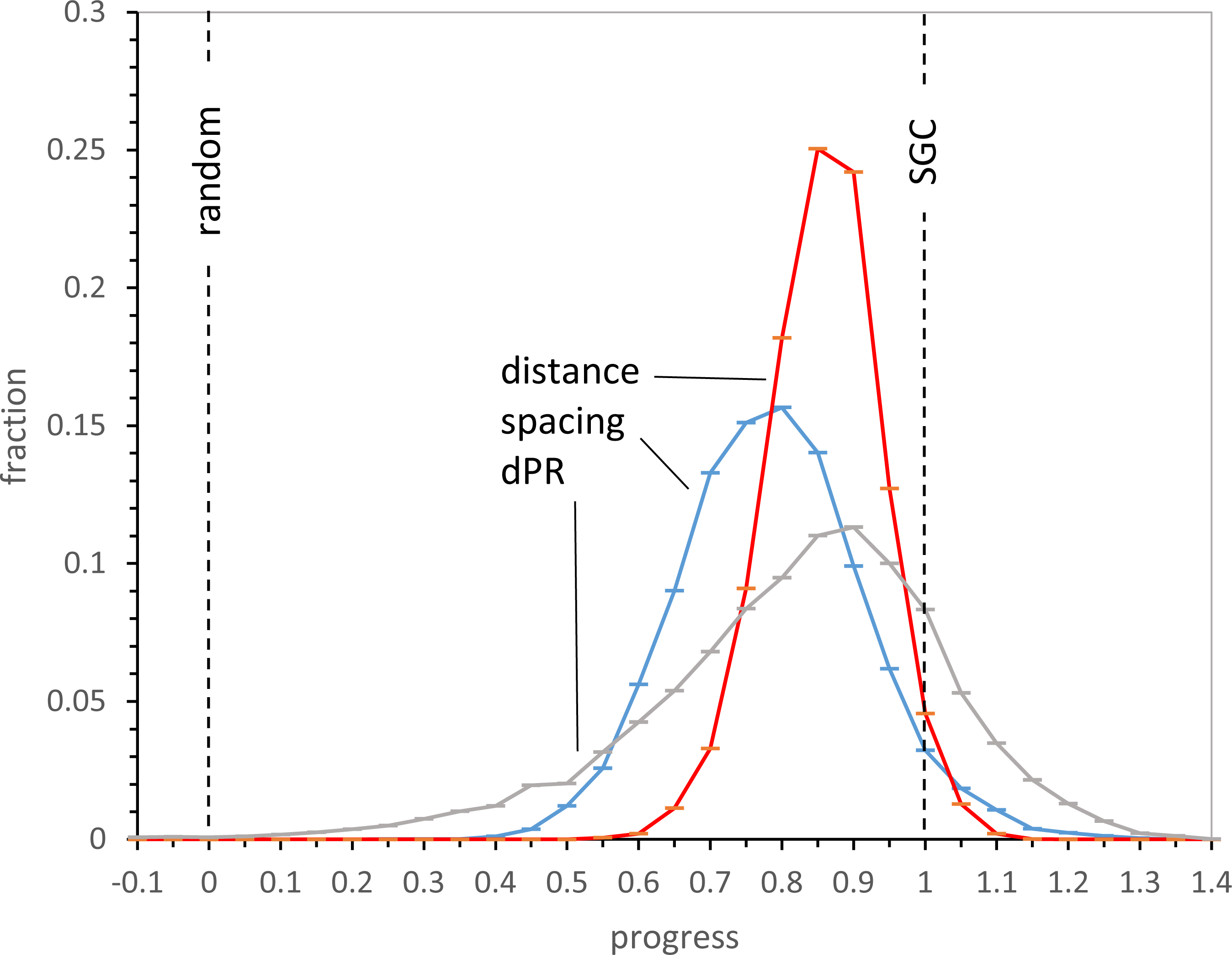
Distributions of progress values with late wobble. Histograms of 10000 late wobble evolutions to 20 encoded functions with Prand = 0.1 and Coevo_PR assignments are shown. Lines mark randomized (“random”) behavior and Standard Genetic Code (“SGC”) behavior. Pinit = 0.6, Pdecay = 0.04, Pmut = 0.04.

**Figure 2C.**
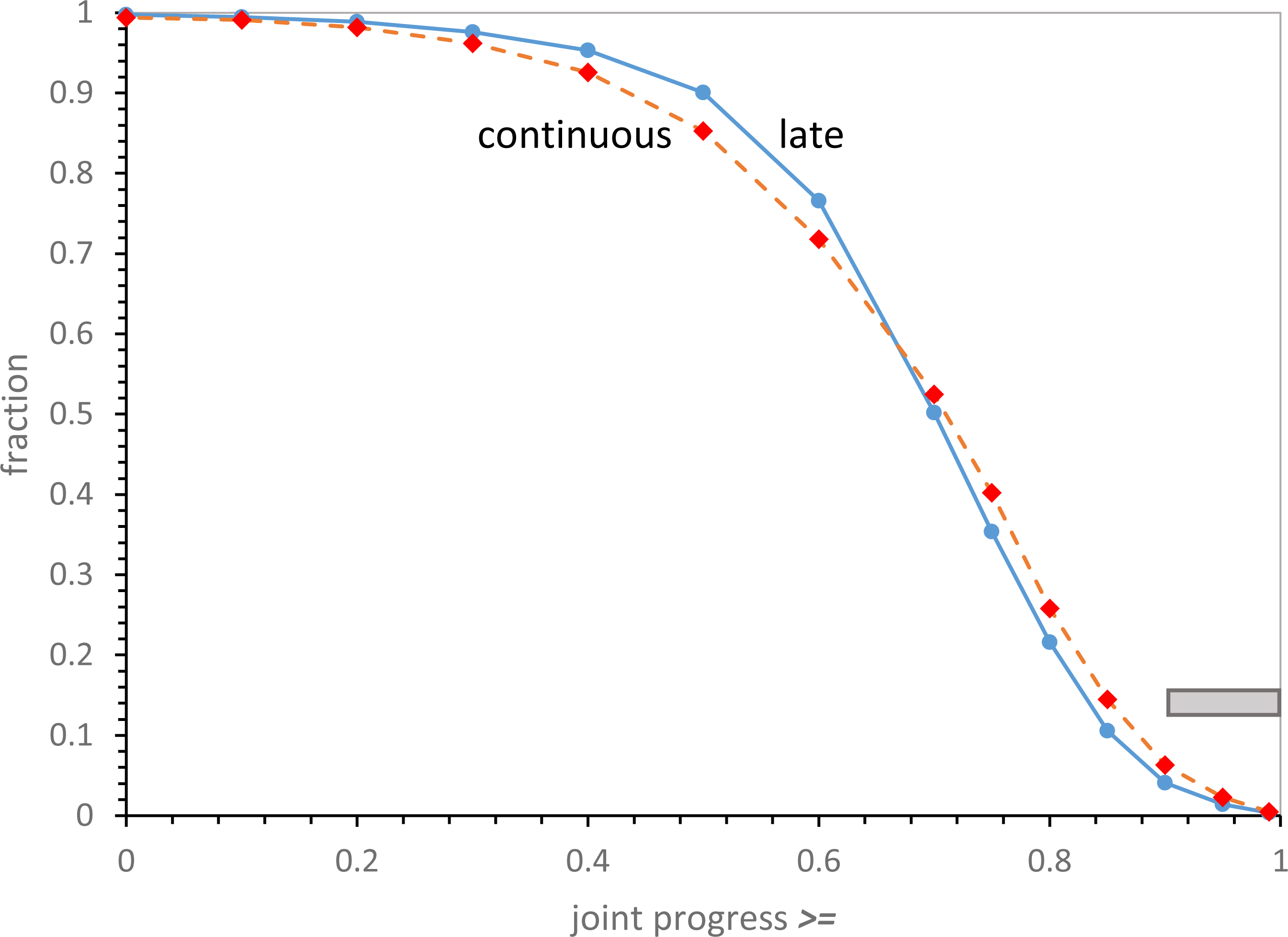
Joint distributions of spacing, distance and dPR progress values for continuous and late wobbling code evolution. Data includes that of Fig. 2B. The “SGC” line indicates all progress values are simultaneously >= 1. Gray bar marks the range for SGC-proximal coding tables; joint progress >= 0.9. 1000 continuous wobbling (to 175 passages) and 10000 late wobbling (to 20 encoded functions) evolutions were employed, with Coevo_PR assignments and Prand = 0.1, Pinit = 0.6, Pdecay = 0.04, Pmut = 0.04, Pwob = 0.5.

**Figure 3.**
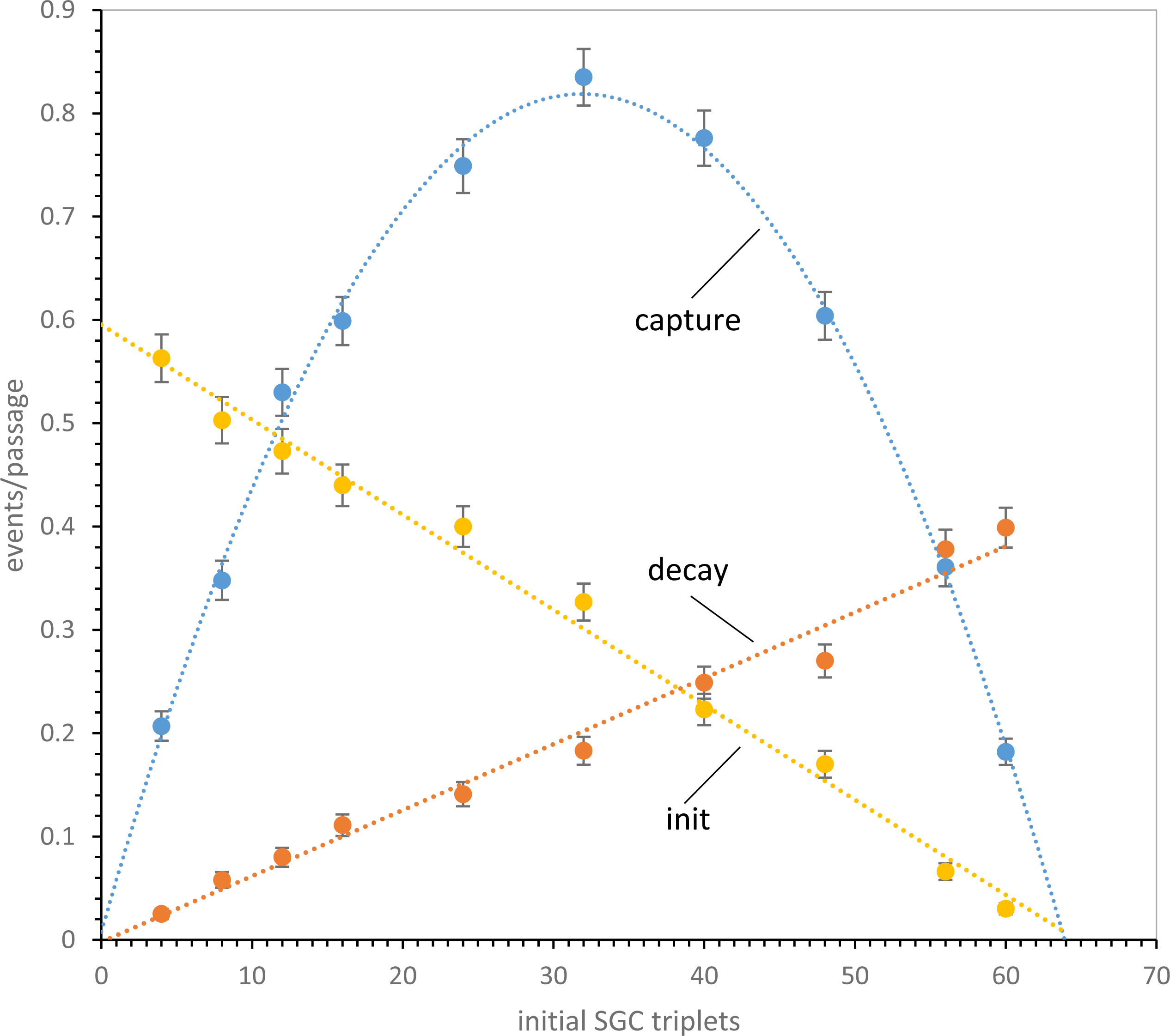
Observed rates of initiation, decay and mutational capture in coding tables with varied numbers of random SGC triplets. Measured mean rates (events/passage ± sem) are shown for 1000 randomly composed non-wobbling tables. Fitted least squares dotted lines portray expected kinetic behavior for initiation, decay, and mutational capture. Decays and captures have been enlarged 10-fold to visualize them on the same scale as initiations. Pmut = 0.04, Pdecay = 0.04, Pinit = 0.6, but Prand = 1.0.

**Figure 4A.**
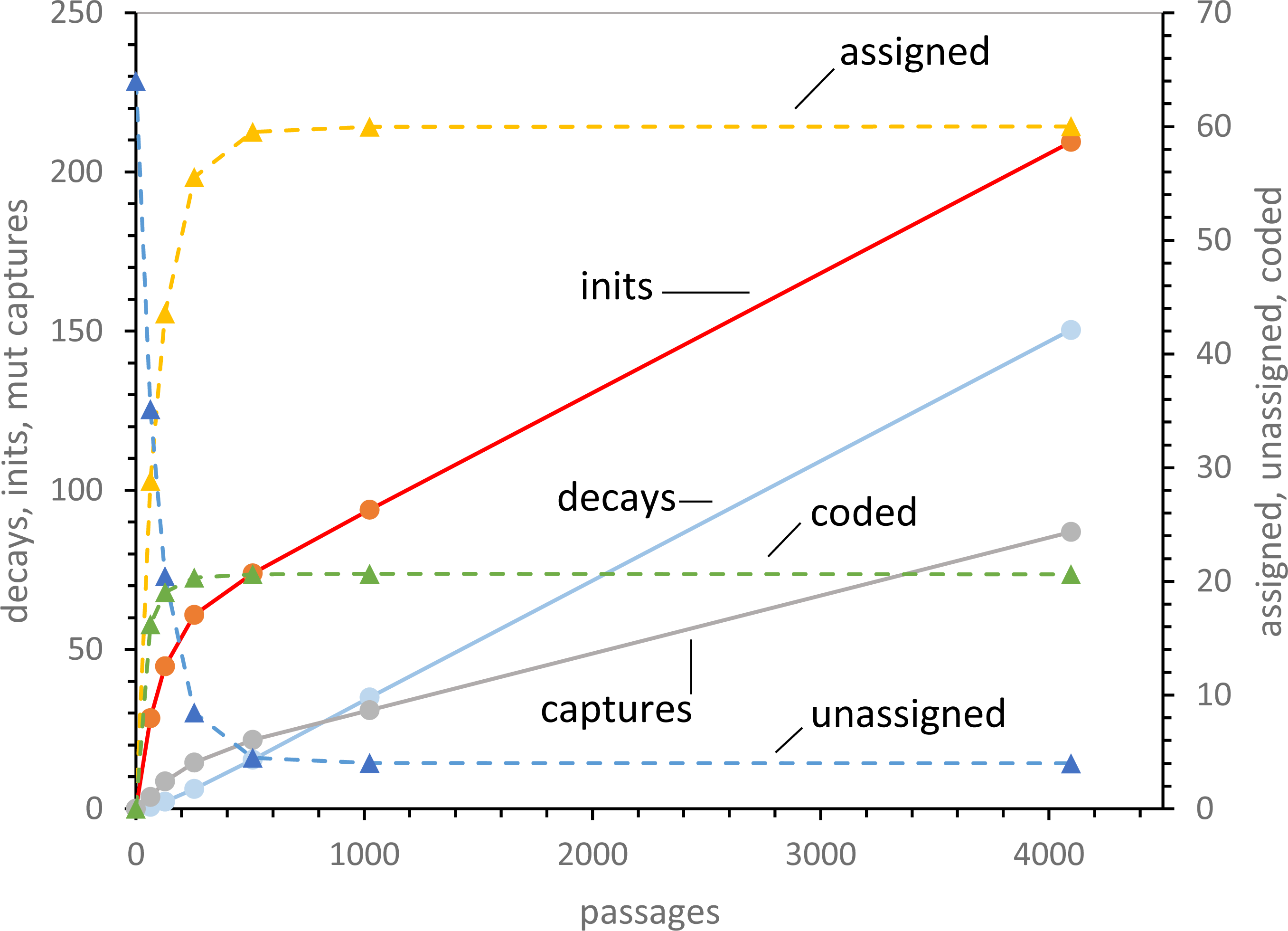
Kinetic evolution of non-wobbling coding tables. Means are shown for 1000 coding tables using random initial assignment and 0 ± 2 PR mutational capture through 4096 passages. Initiations, decays and mutational captures are plotted, with triplets assigned, triplets unassigned and functions coded on a secondary axis (rightward). Transition probabilities as in Fig. 3.

**Figure 4B.**
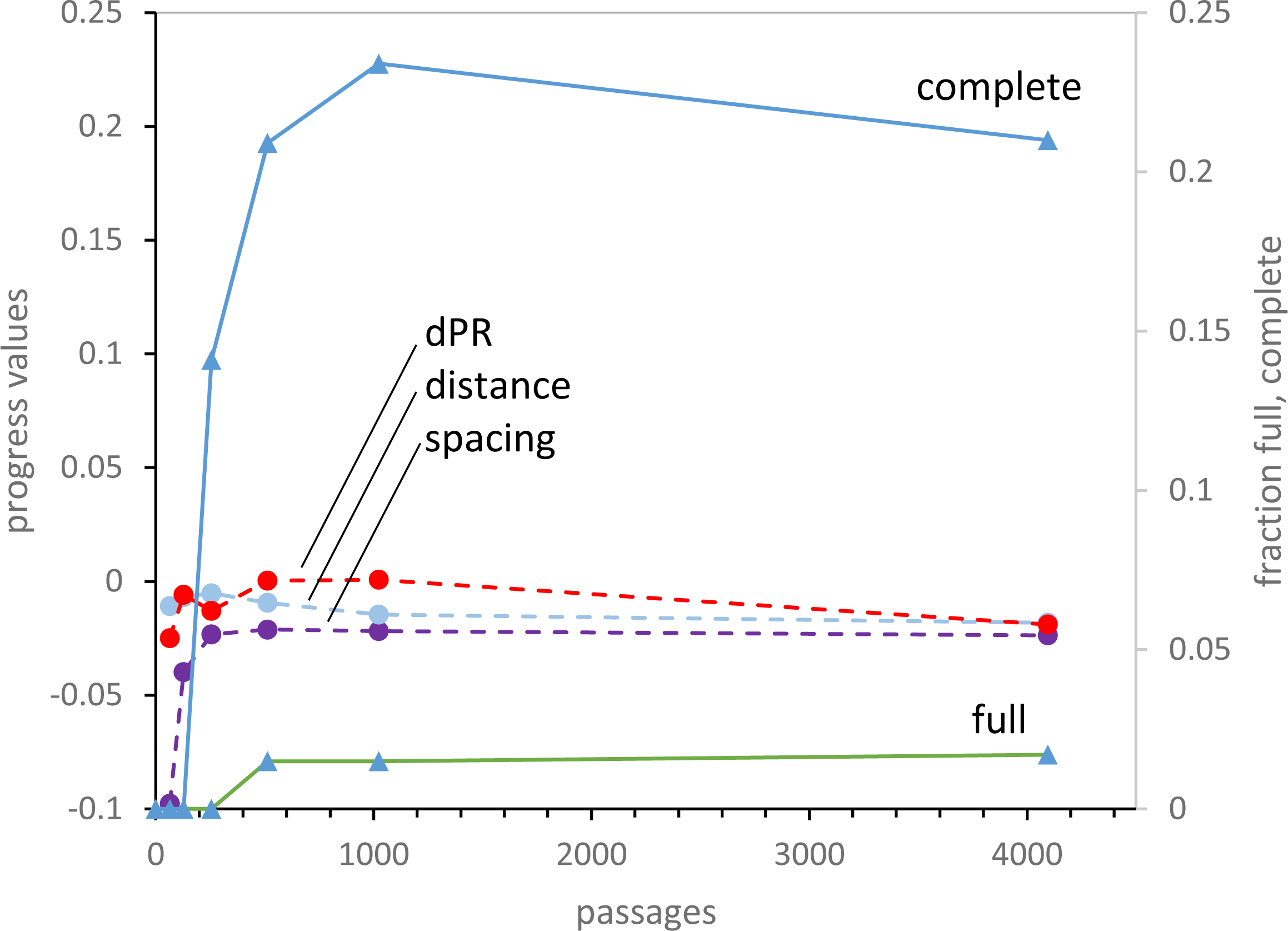
Progress and finished non-wobbling coding tables. Means of 1000 spacing, distance and dPR progress values during the 4096 passages of Fig. 4A are plotted on the left ordinate, and the fraction of full and complete coding tables are on the right ordinate.

**Figure 5A.**
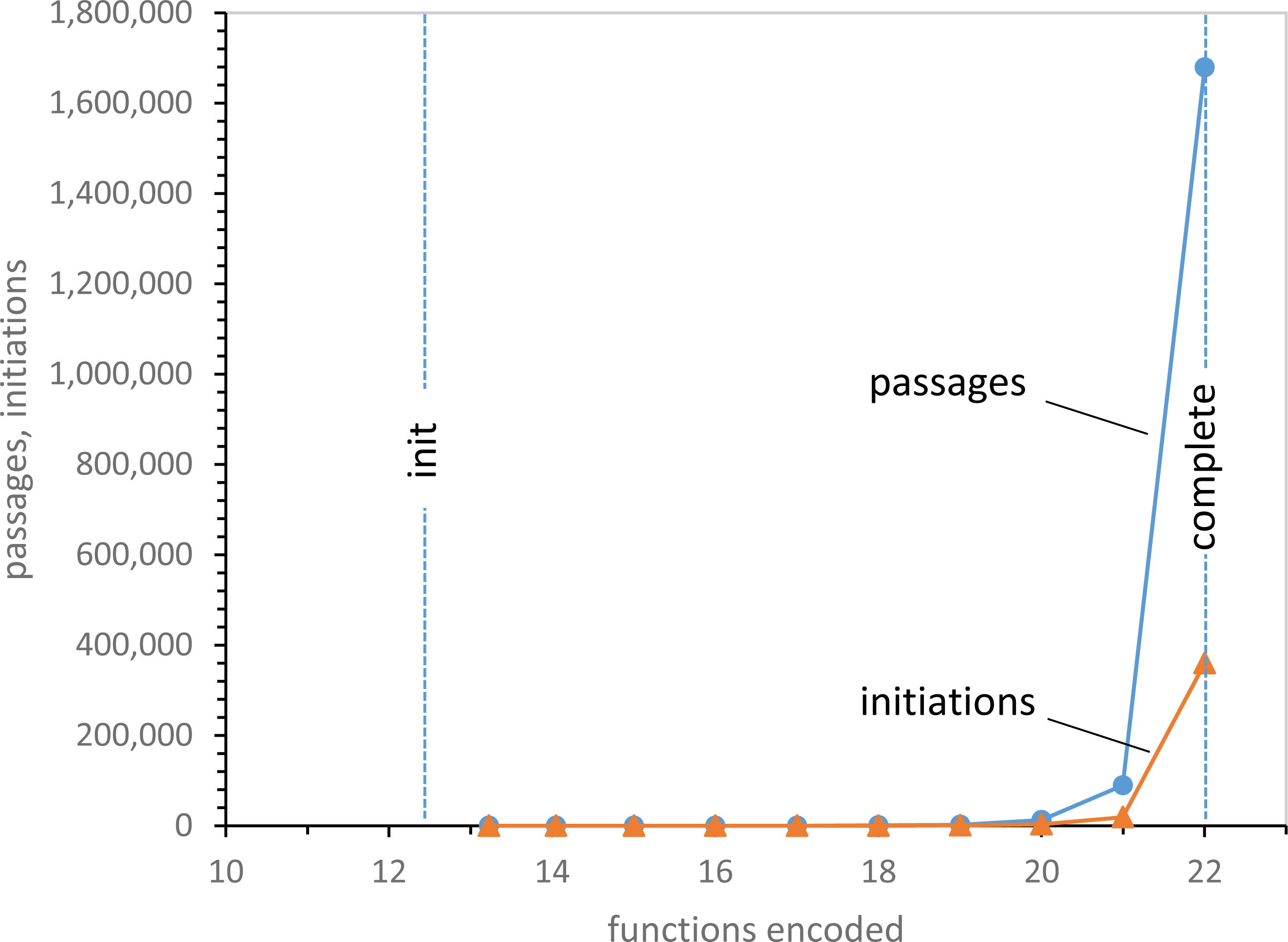
Mean passages and initiations required for random wobble encoding of varied numbers of functions, after 16 random SGC wobble initiations. 1000 evolutions using random wobble coding and Coevo_PR mutational capture were stopped at numbers of encoded functions on the abscissa. Lines labeled “init” are just after 16 initial random SGC triplets are chosen, but before further evolution. The “complete” line marks acquisition of 22 encoded functions. Pmut = 0.04, Pdecay = 0.04, Pinit = 0.6, Pwob = 0.5, Prand = 1.0.

**Figure 5B.**
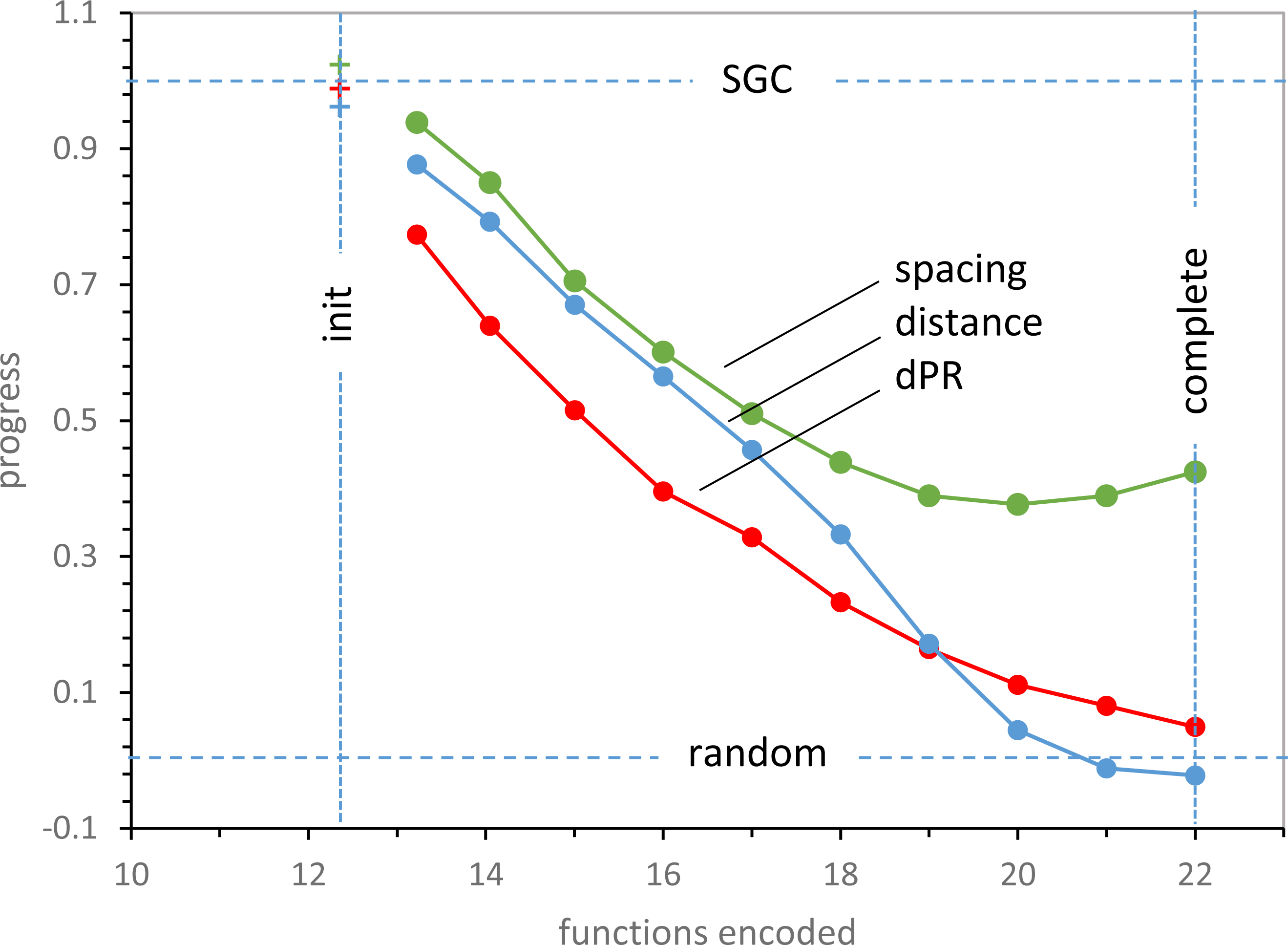
Mean spacing, distance and dPR progress values for random wobble encoding after 16 random SGC wobble initiations. Data from calculations in Fig. 5A. **“**Init” and “complete” lines as in Fig. 5A. “SGC” line indicates SGC-like progress values, “random” line indicates progress for randomized assignments.

**Figure 5C.**
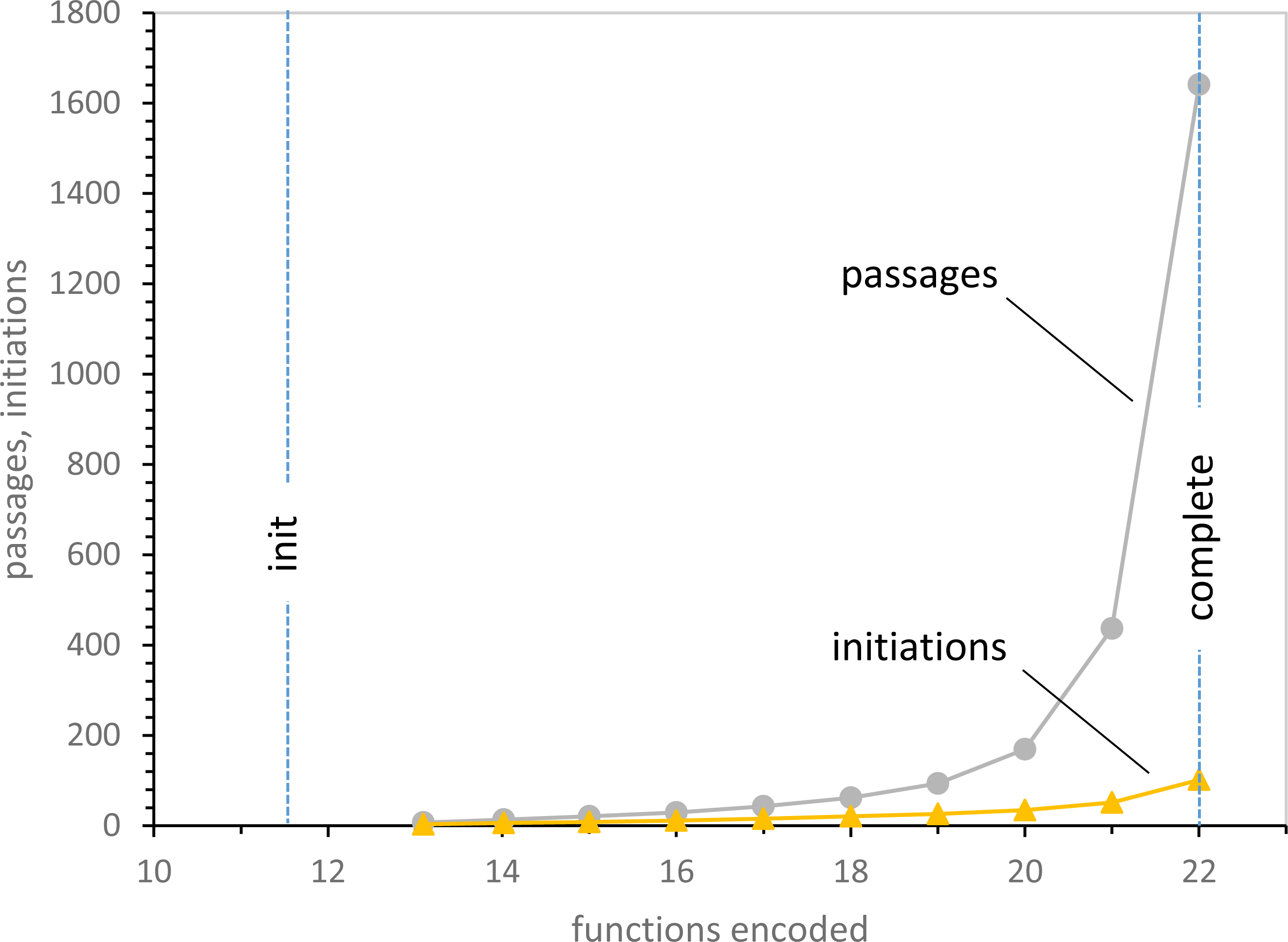
Mean passages and initiations required for random non-wobble encoding, after 16 initial random SGC non-wobble initiations. 1000 evolutions using random wobble coding and Coevo_PR mutational capture were stopped at numbers of encoded functions on the abscissa. Lines labeled “init” indicate positions just after initial 16 random SGC triplets are chosen. Lines labeled “complete” indicate acquisition of 22 encoded functions. Transition probabilities as in Fig. 5A.

**Figure 5D.**
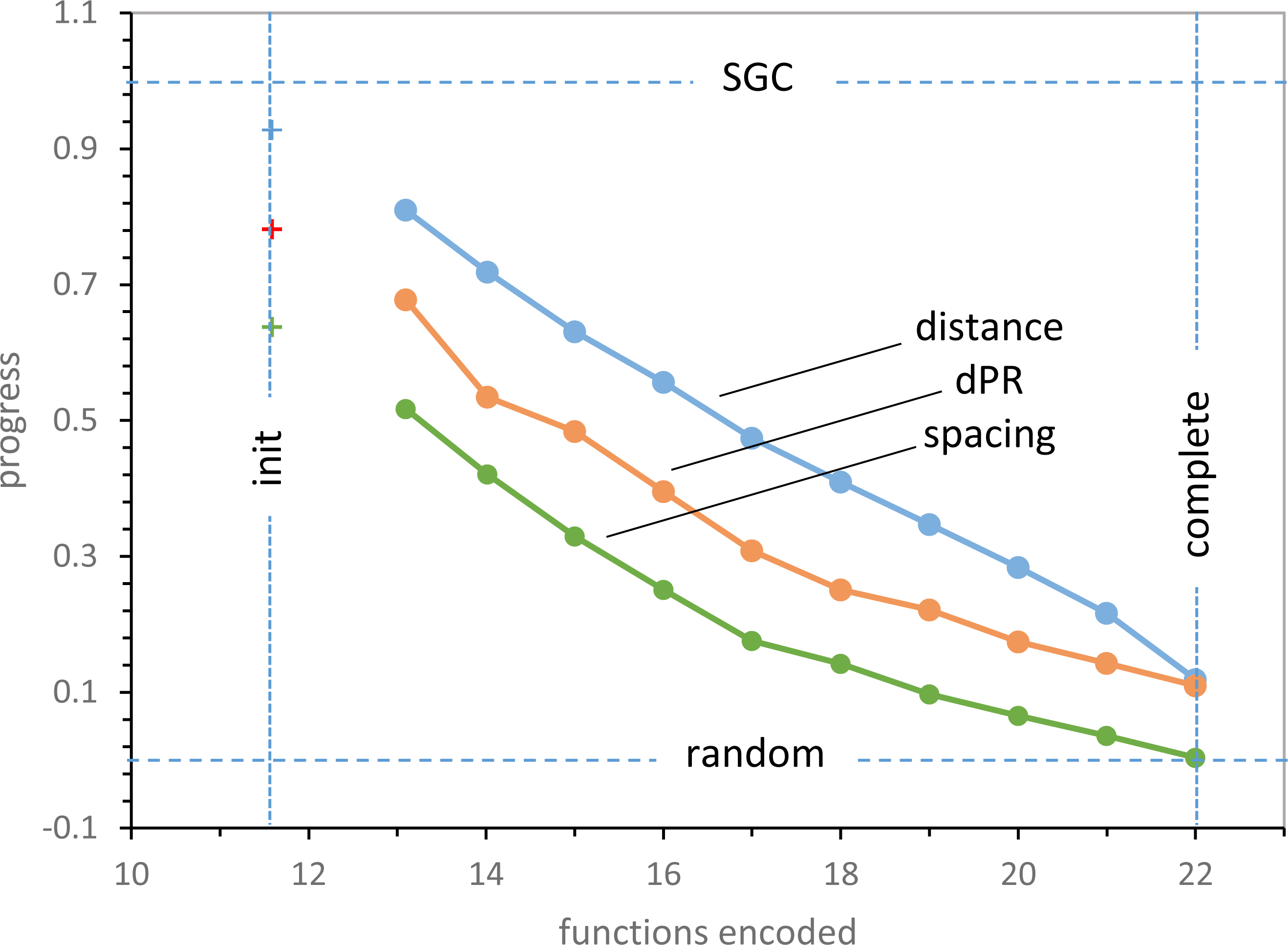
Mean spacing, distance and dPR progress values for random non-wobble encoding after 16 initial random SGC non-wobble initiations. Data from calculations in Fig. 5C. Labeling as in Fig. 5B.

**Figure 6A.**
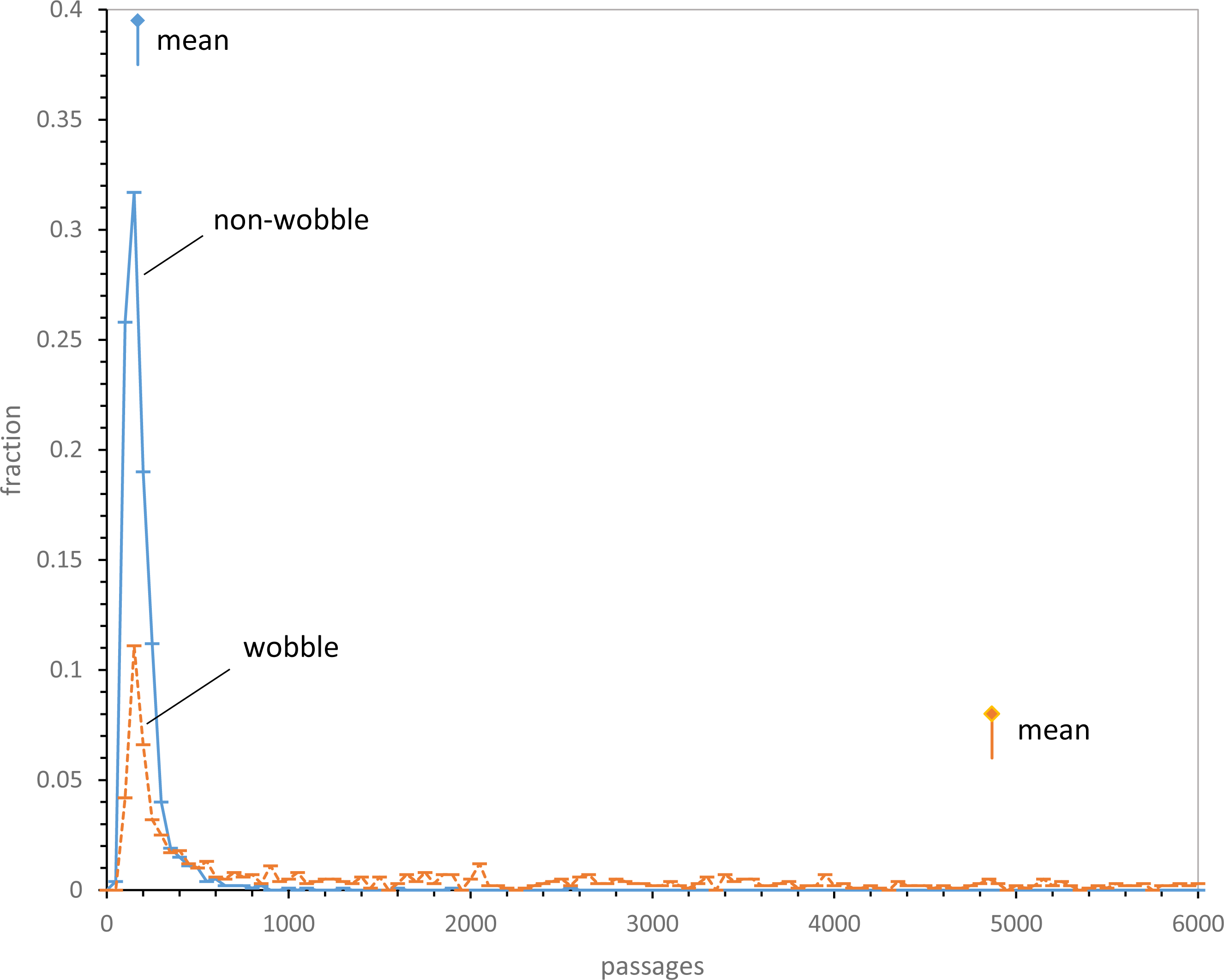
Fraction of near-complete 20-function codes after varied durations. 1000 evolutions with Coevo_PR captures and 10% random assignment were applied to wobble and non-wobble coding. Probabilities except for Prand = 0.1, as in Fig. 5A. Signpost symbols are distribution means. All non-wobble evolutions are plotted, but 0.216 of wobble evolutions required longer durations and are off-scale to the right.

**Figure 6B.**
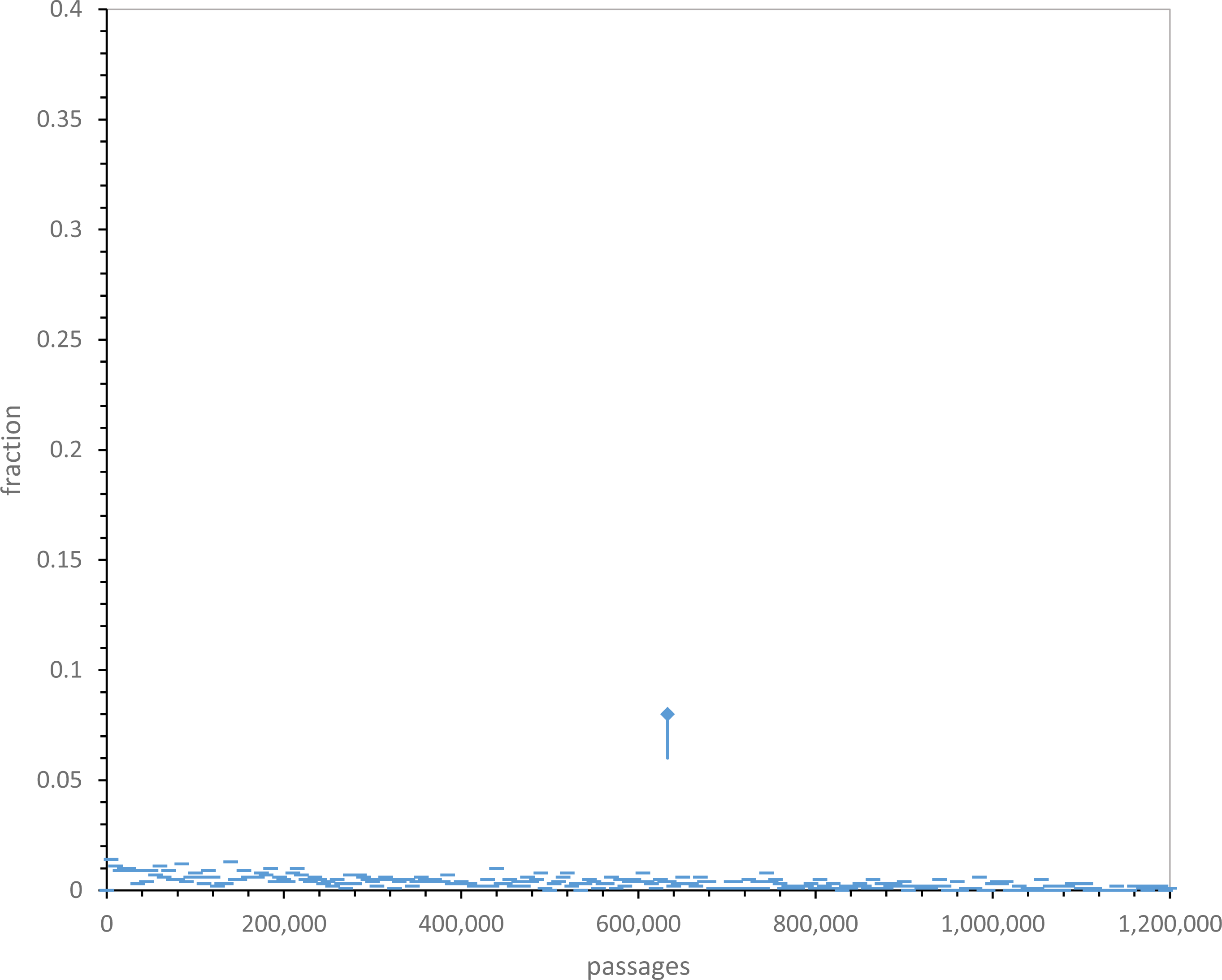
Fraction of complete 22-function wobble codes at varied durations. 1000 evolutions with Coevo_PR assignments and Prand = 0.1 were used. Other probabilities as for Fig. 6A. Signpost symbol marks the mean. 0.142 of complete wobble evolutions required longer durations, off-scale to the right.

**Figure 7A.**
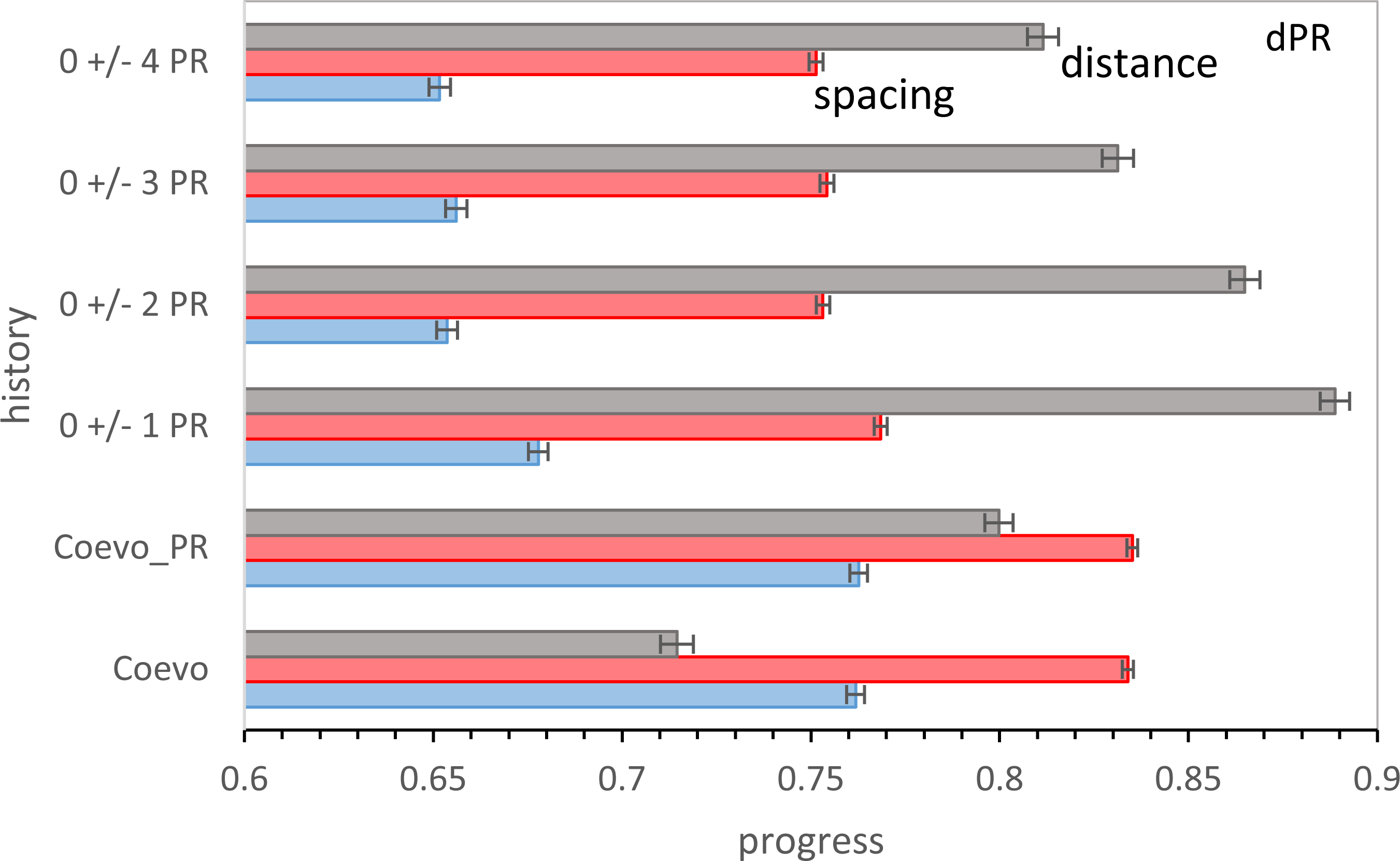
Progress for 6 mutational assignment histories using late wobble. 3000 evolutions were averaged for non-wobble encoding to 20 functions, then all possible late wobbles. Bars are sem’s. Pmut = 0.04, Pdecay = 0.04, Pinit = 0.6, Prand = 0.1.

**Figure 7B.**
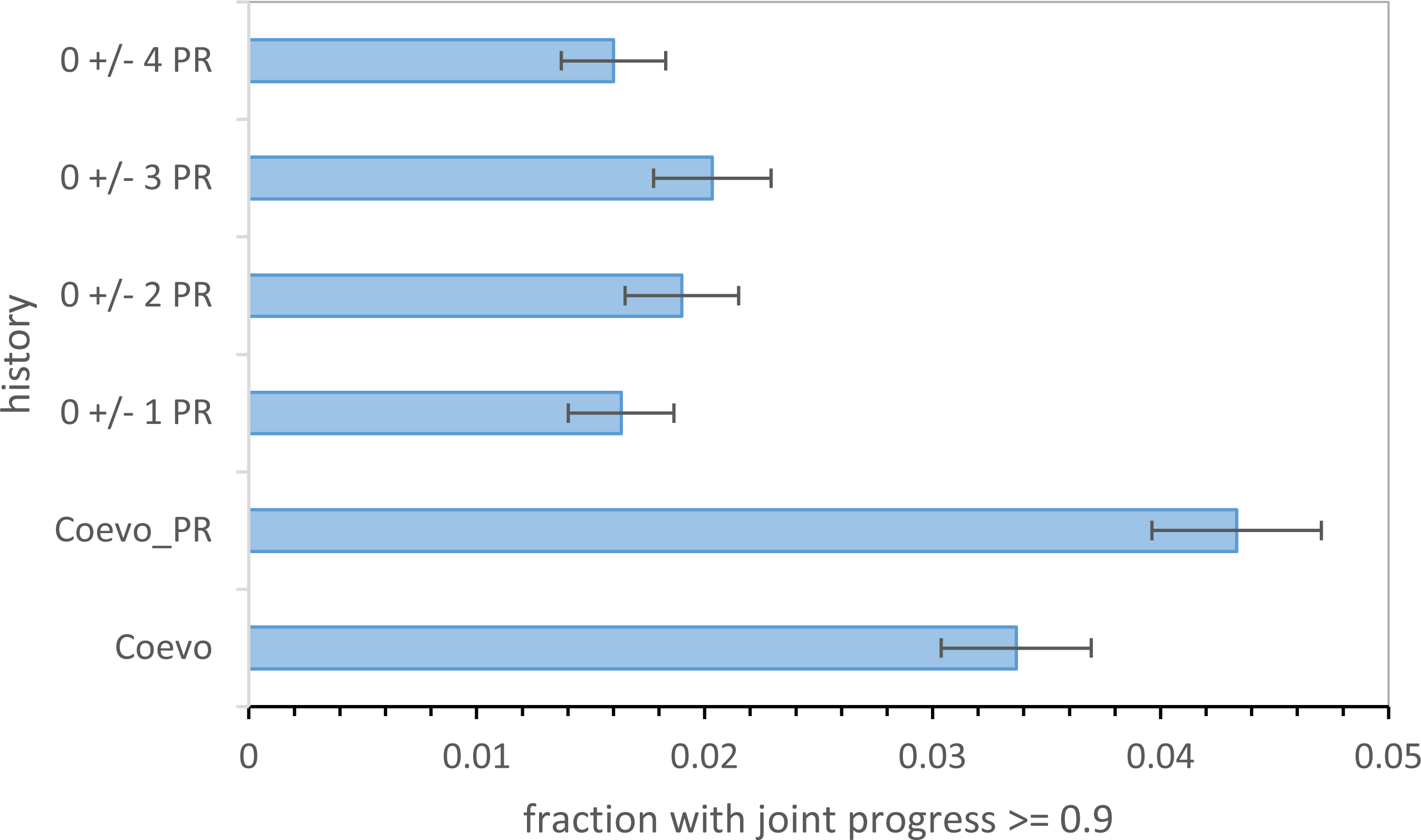
Fraction of evolutions with joint progress for 6 mutational assignment histories using late wobble. Spacing, distance and dPR progress values from 3000 late wobble evolutions were used, and joint progress is plotted with standard errors. Data are from Figure 7A.

**Figure 7C.**
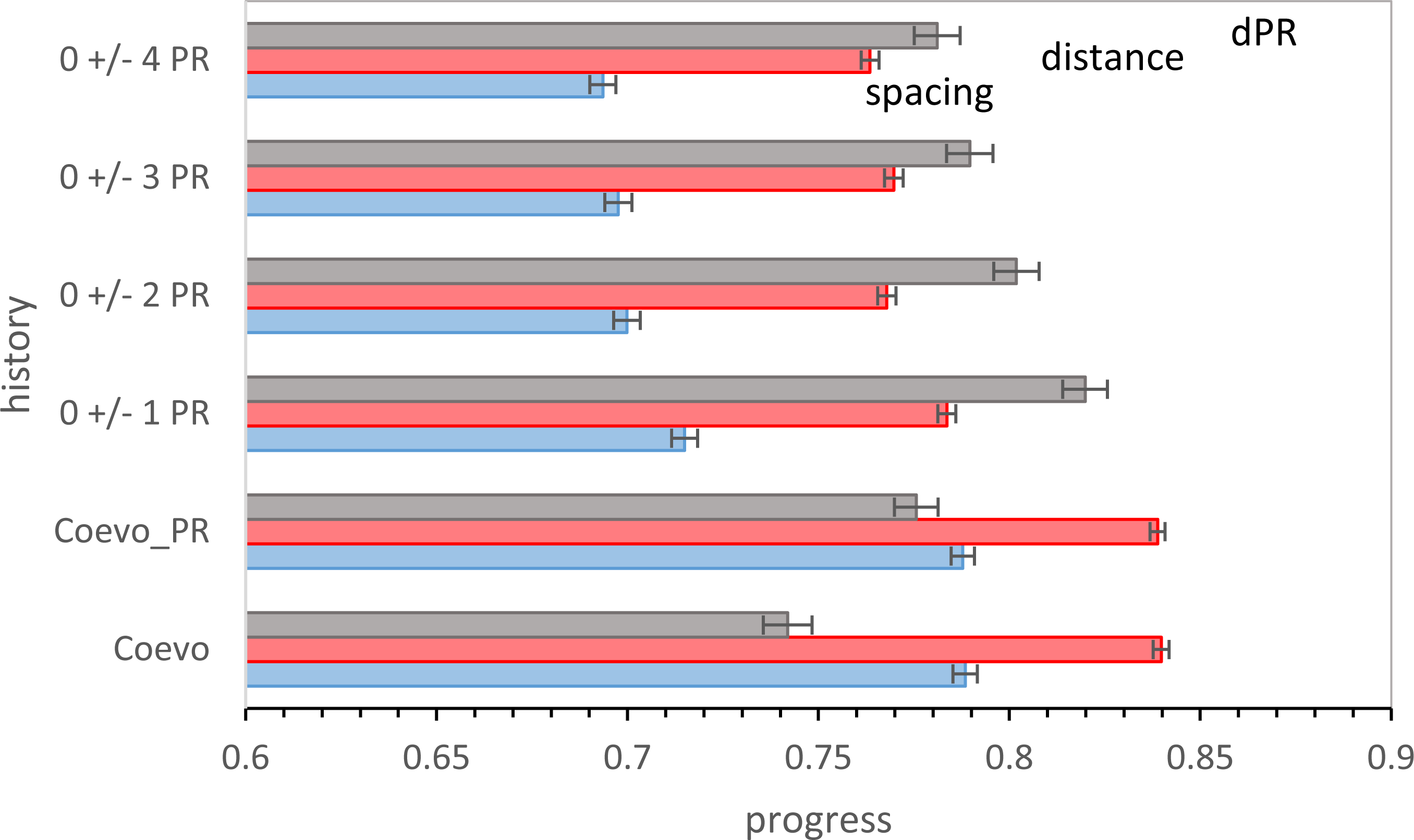
Progress for 6 mutational assignment histories using continuous wobble. Spacing, distance and dPR progress values from 2000 continuous wobble evolutions’were averaged, and plotted with standard errors. Pmut = 0.04, Pdecay = 0.04, Pinit = 0.6, Pwob = 0.5, Prand = 0.1.

**Figure 7D.**
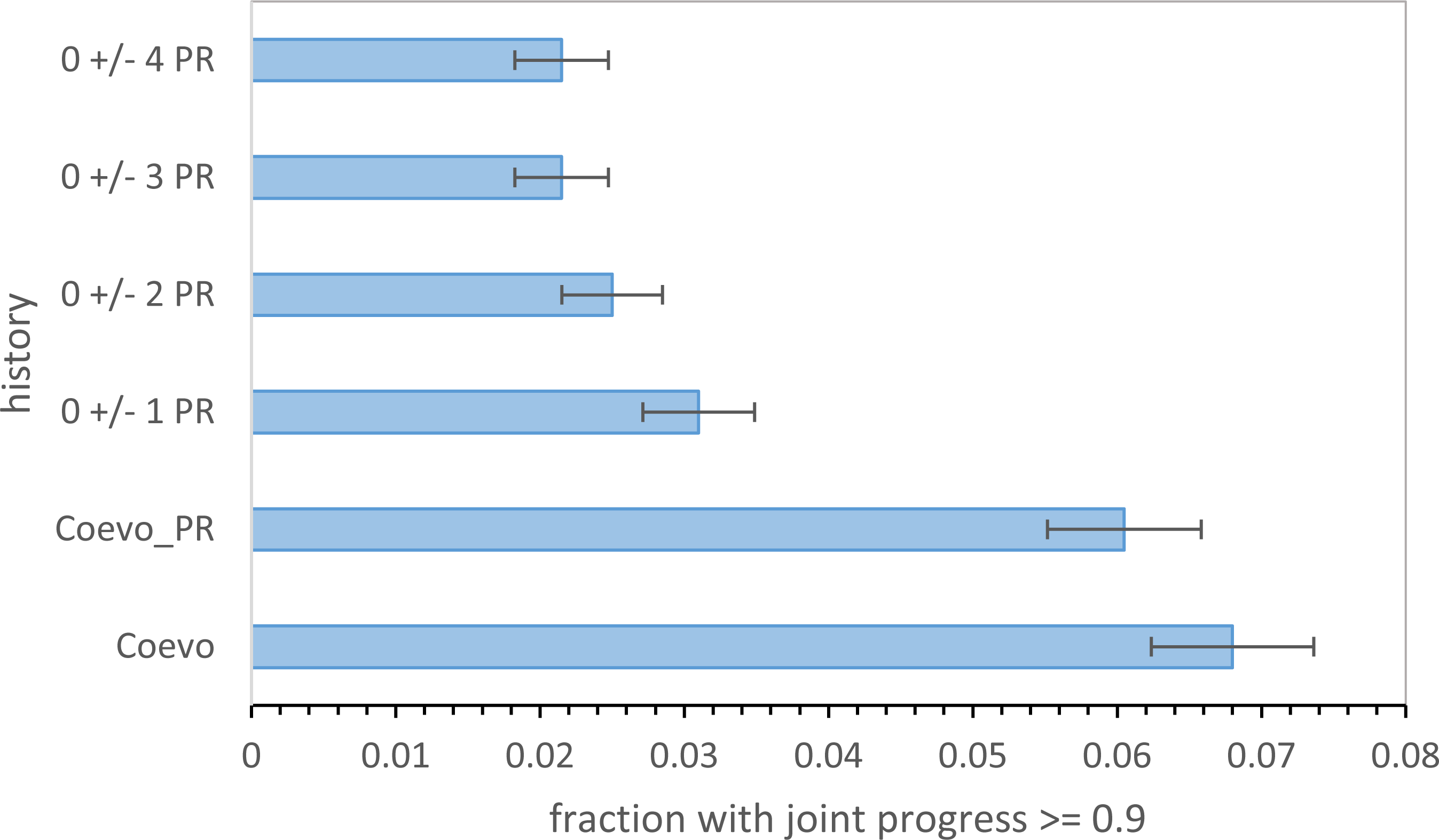
Fraction of evolutions with joint progress for 6 mutational assignment histories using continuous wobble. Fraction of 2000 continuous wobble evolutions with spacing, distance and dPR progress simultaneously >= 0.9 were plotted with standard errors. Data are from Figure 7C.

**Figure 8A.**
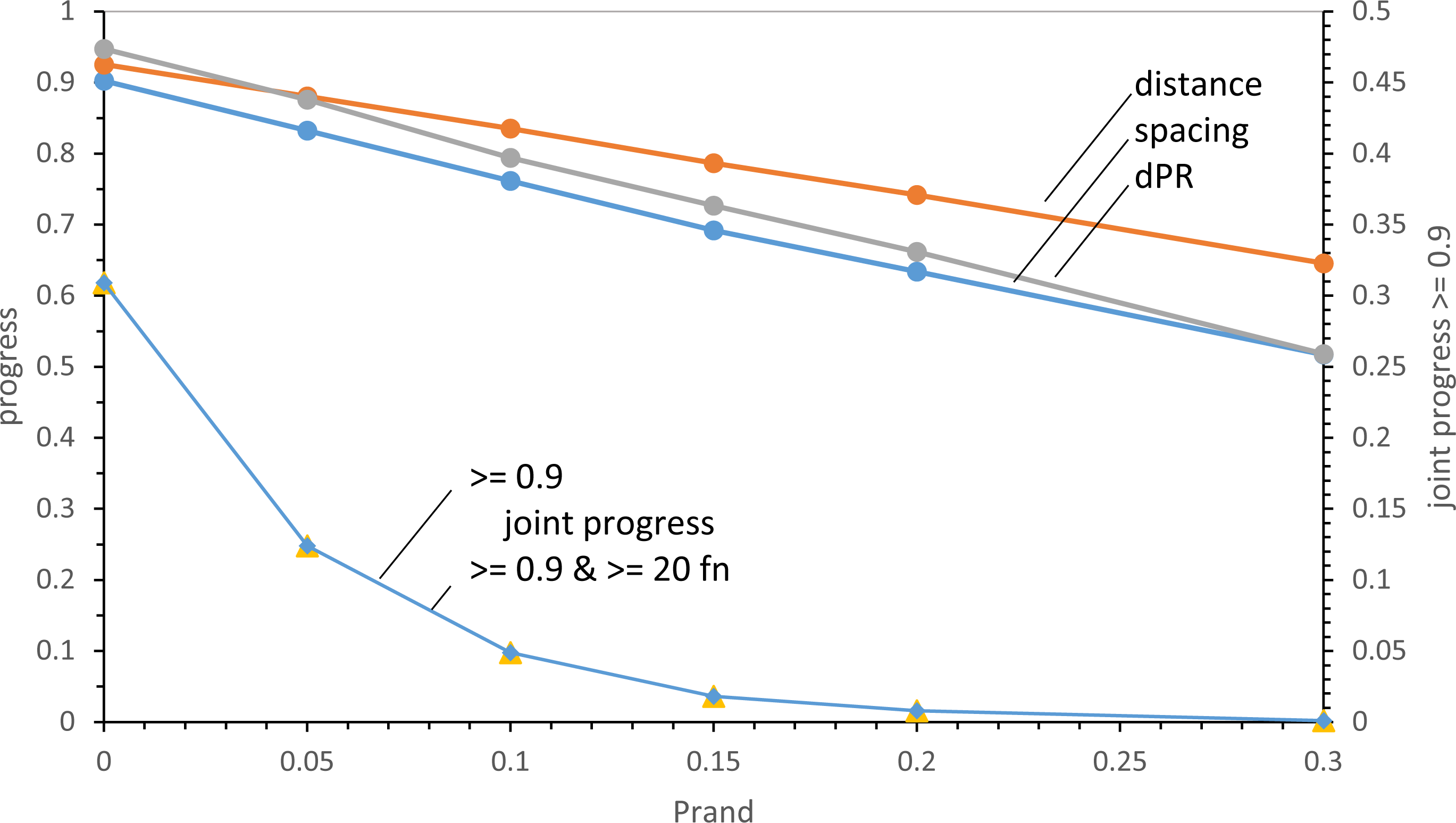
Effects of random assignments on late wobbling code order. 1000 evolutions to 20 encoded functions with Pmut = 0.04, Pdecay = 0.04, Pinit = 0.60 and varied Prand were used. Progress values plotted on left ordinate, fraction of evolutions with joint progress on right.

**Figure 8B.**
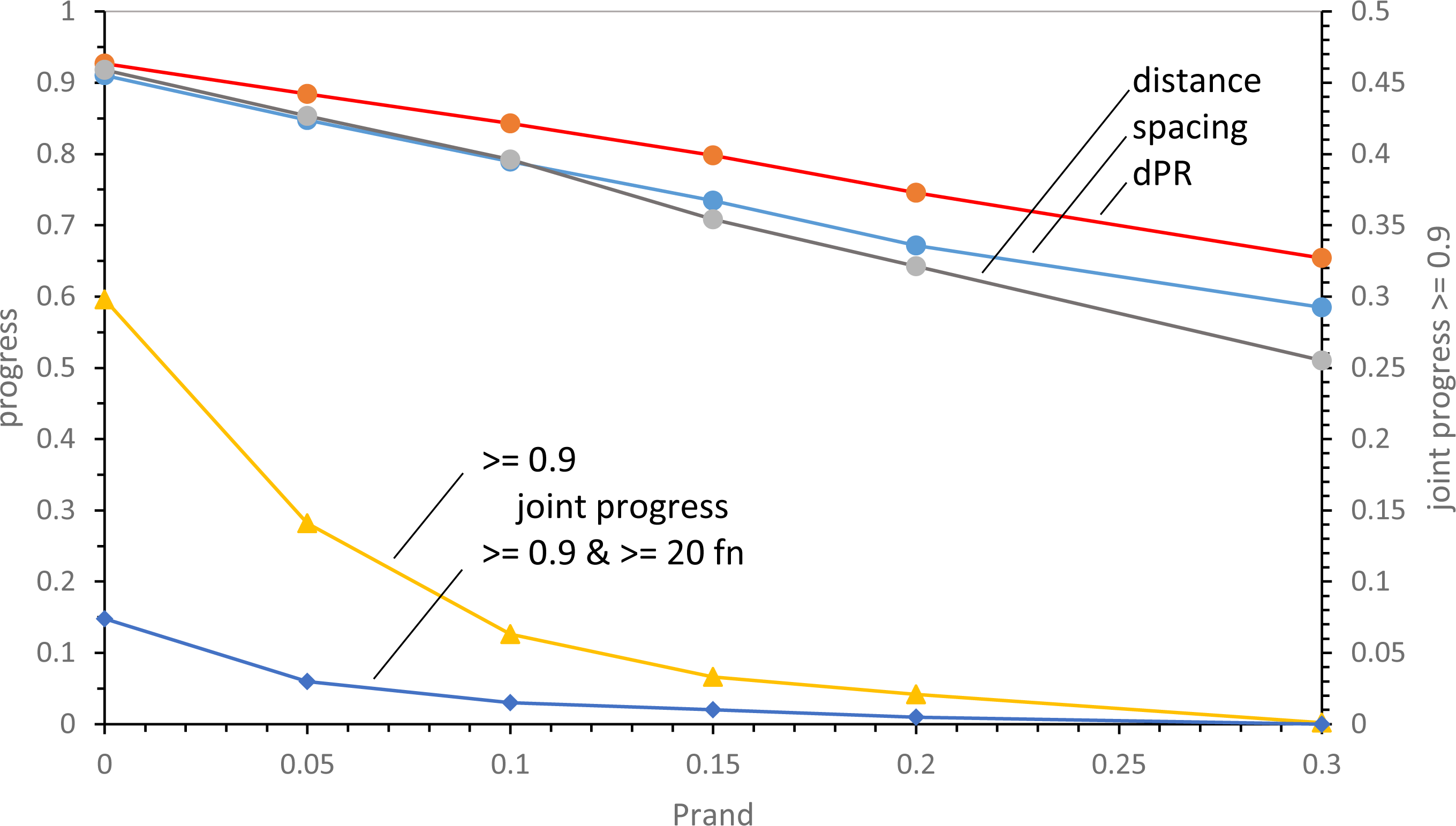
Effects of random assignments on continuous wobbling code order. 1000 evolutions to 175 passages with Pwob= 0.5 and otherwise, the same probabilities and plotting as in Fig. 8A.

**Figure 9A.**
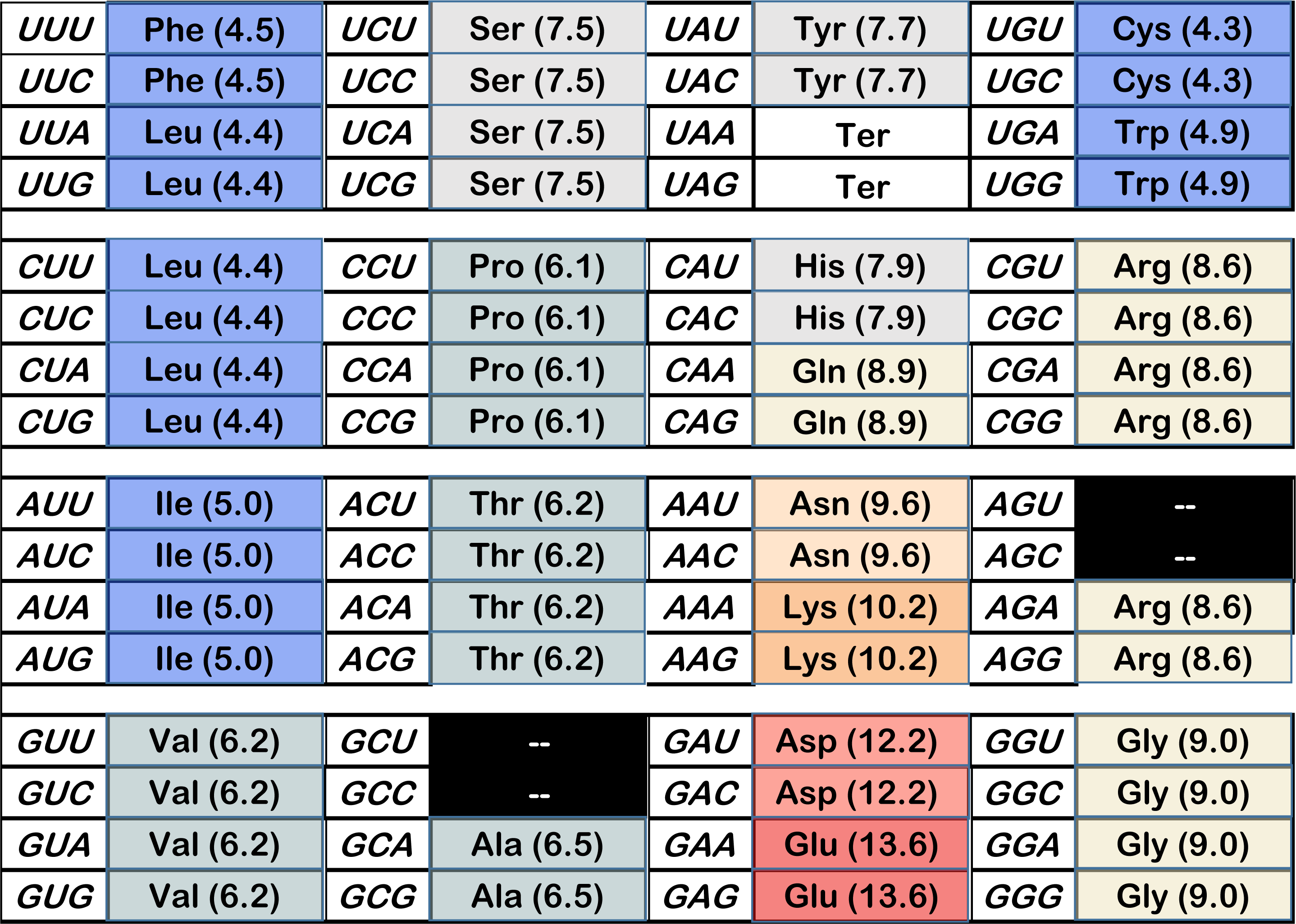
Arguably the best late wobble coding table observed in 600. spacing progress, 1.129, distance, 1.048; dPR, 0.880. Frequency approximately 1 of 600: number 294/600: relation to SGC – 2 altered assignments (AUG Ile and UGA Trp), 4 unassigned, but SGC chemical ordering (Fig. 1A) fully reproduced.

**Figure 9B.**
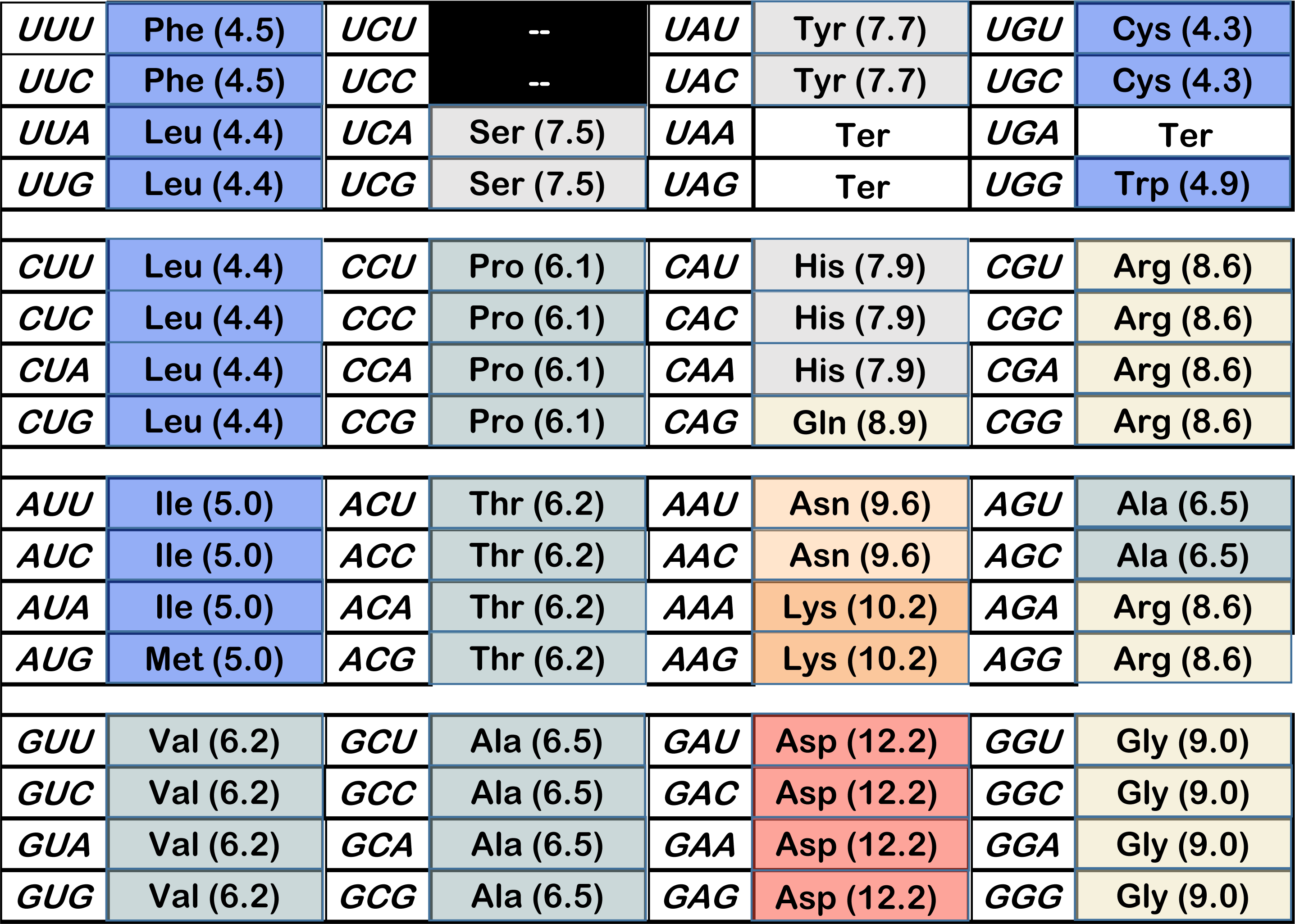
Example, late wobble joint progress ca. 1.0. spacing progress, 0.983; distance, 1.002; dPR, 0.998; frequency of equivalent or better indices = 0.0015: number 23/600: relation to SGC – 3 altered assignments (AGU/C Ala, GAA/G Asp and CAA His), 2 unassigned, but preserves chemical order except for AGY Ser.

**Figure 9C.**
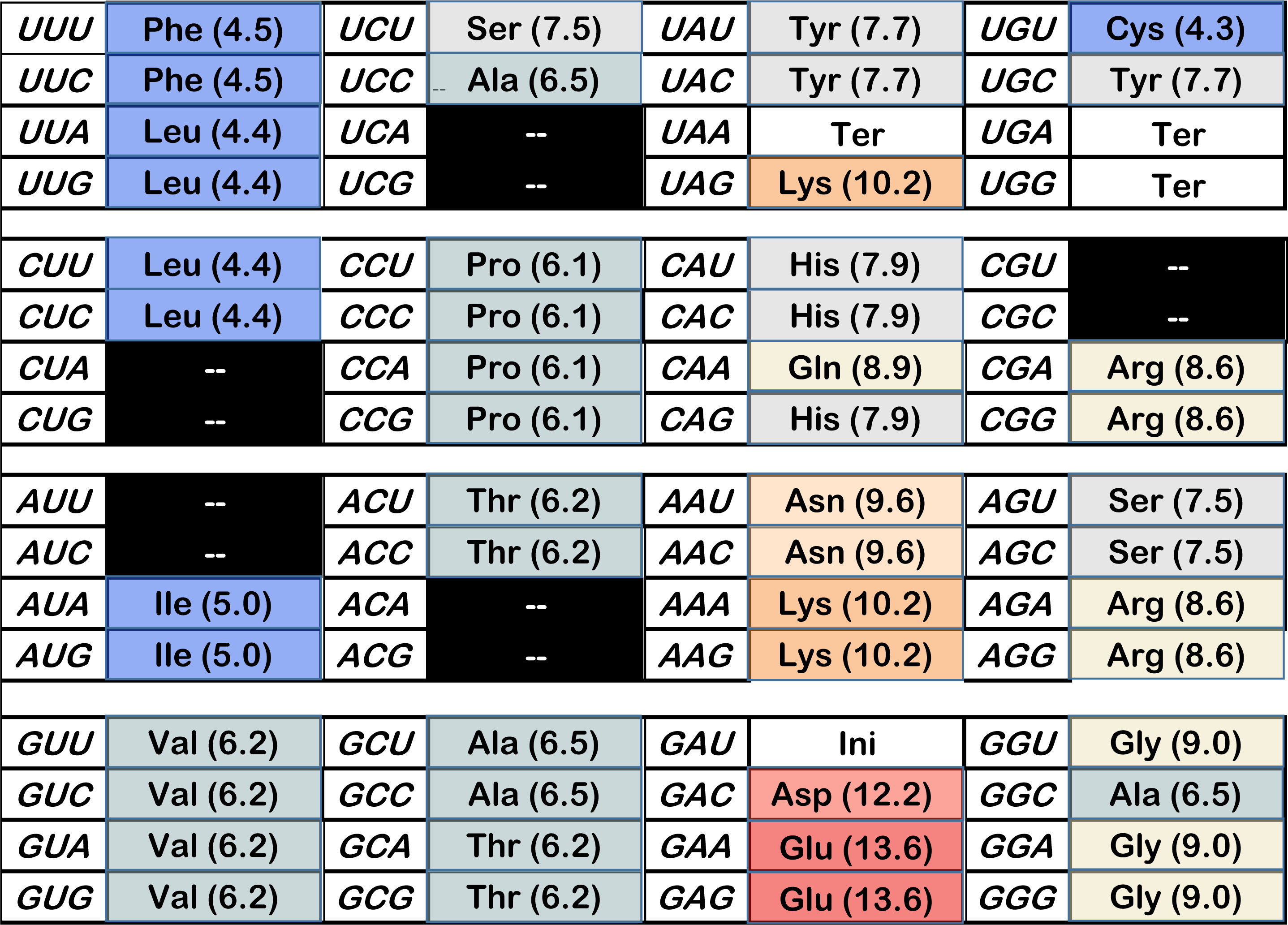
Example, late wobble joint progress ca. 0.95. spacing progress, 0.946, distance, 0.937, dPR, 1.056; frequency of equivalent or better indices, 0.0045: number 6/600: relation to SGC – 10 altered assignments, 10 unassigned, intermediate polar requirement bloc seriously disrupted.

**Figure 9D.**
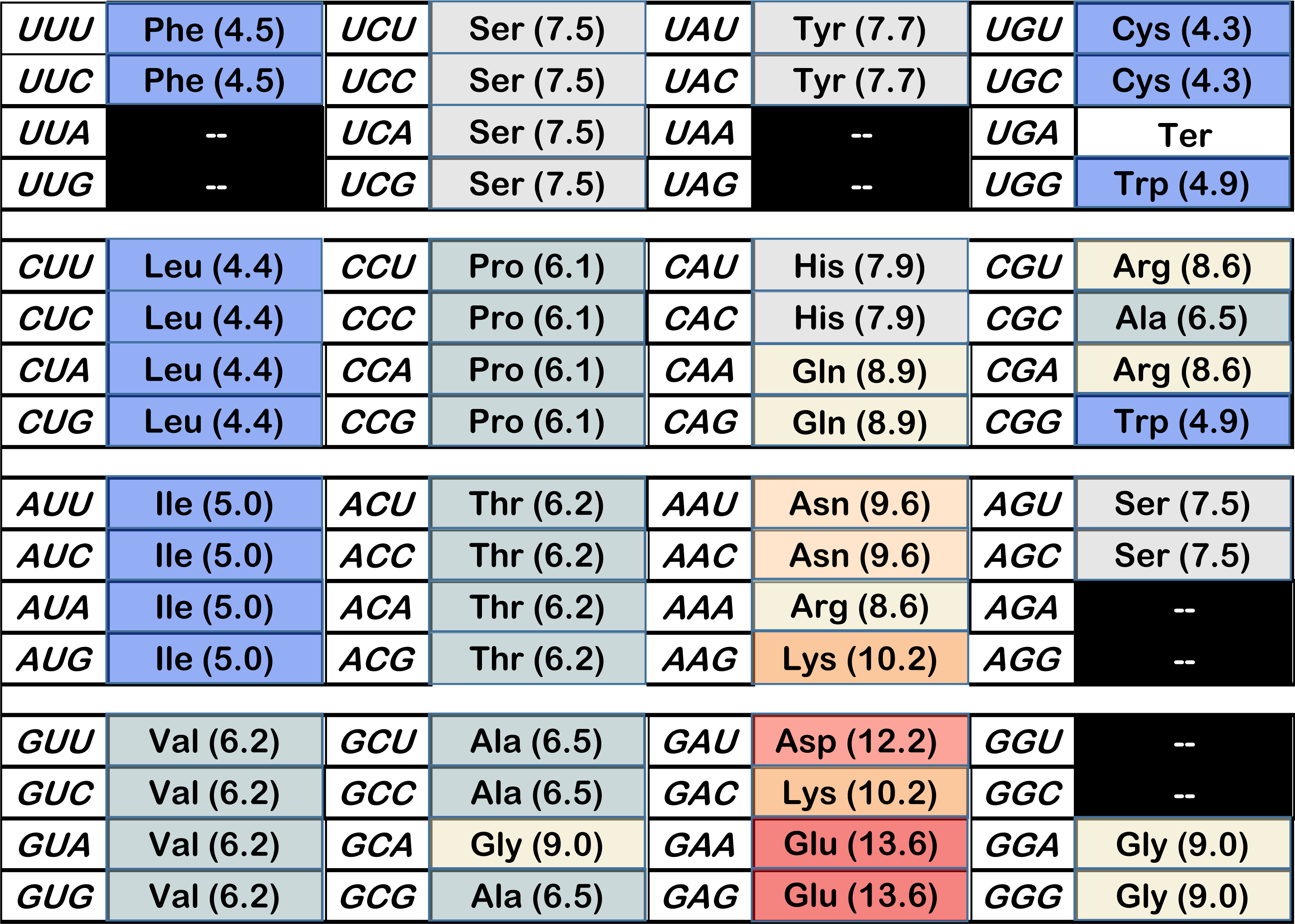
Example, late wobble joint progress ca. 0.9. spacing progress, 0.912, distance, 0.905, dPR, 0.920; frequency of equivalent or better indices, 0.0325: number 286/600: relation to SGC – 6 altered assignments, 8 unassigned, small perturbations in intermediate and very polar blocs.

**Figure 10A.**
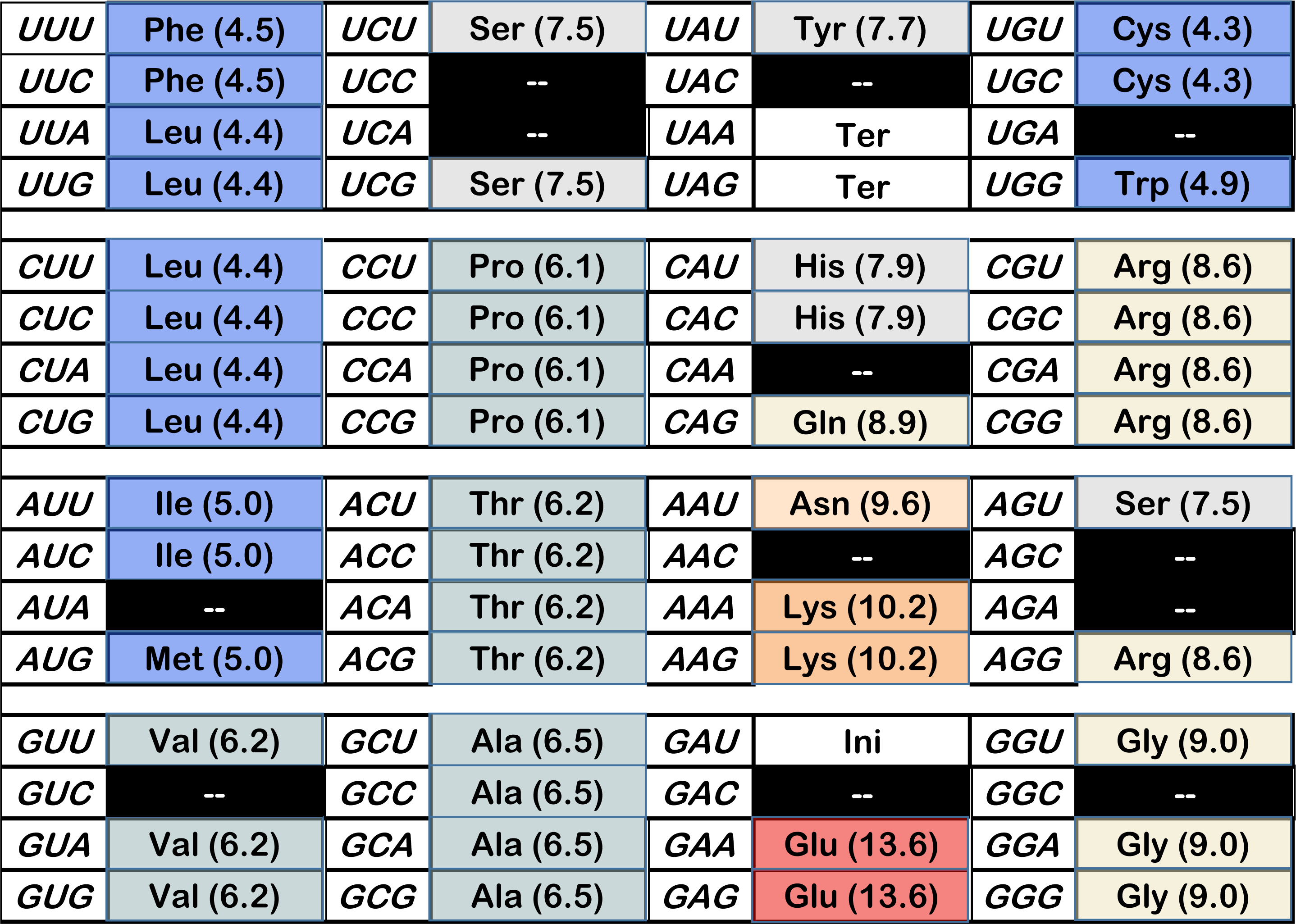
Arguably the best continuous wobble coding table of 600. spacing progress, 1.072, distance, 1.031; dPR, 1.114. Frequency approximately 1 of 600: number 549/600: relation to SGC – 1 altered assignment (GAU Ini), 12 unassigned, but reproduces SGC chemical ordering fully (compare Fig. 1A). The only 21-function coding table among examples.

**Figure 10B.**
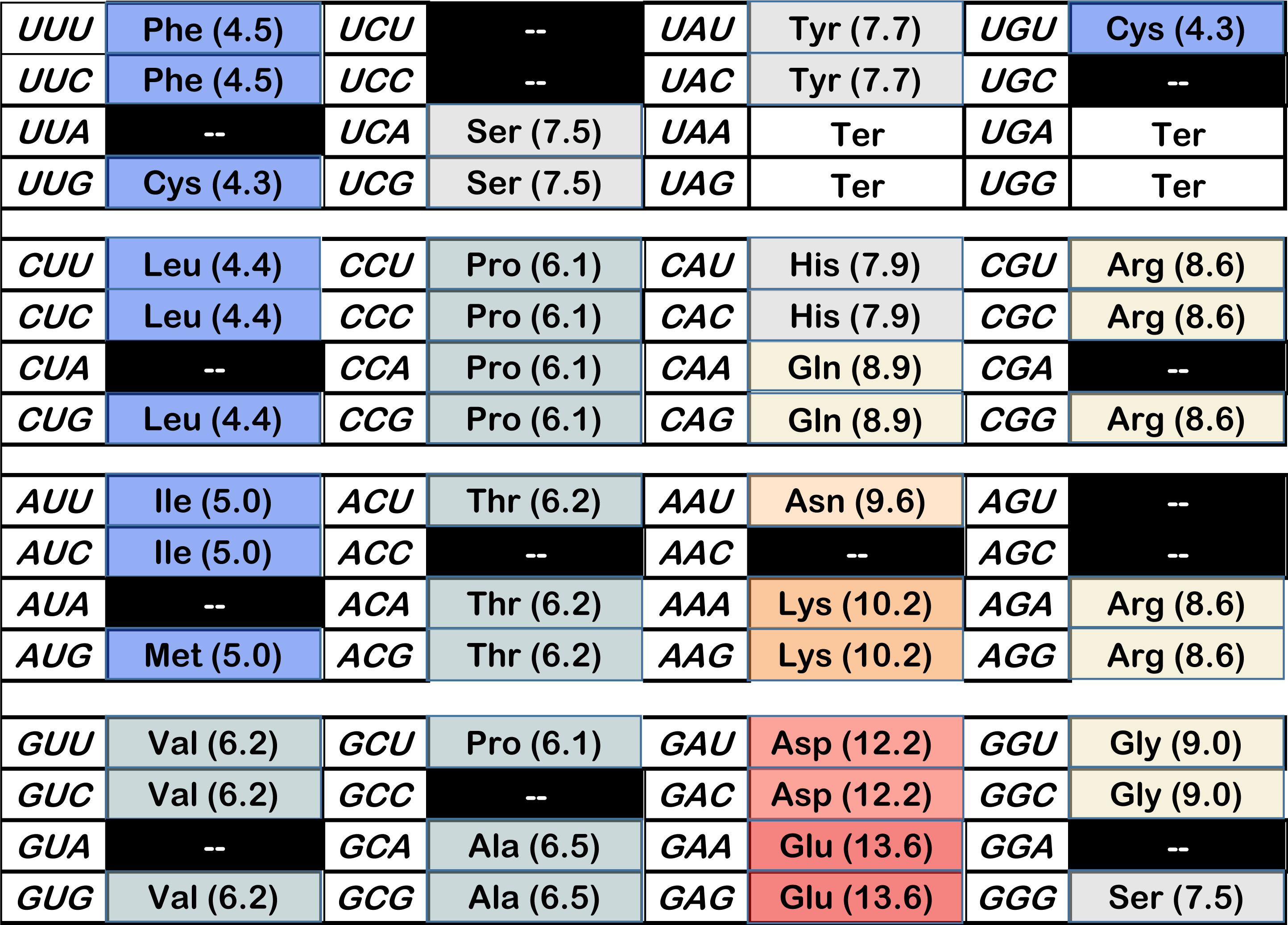
Example, continuous wobble joint progress ca. 1.0. spacing progress, 0.983; distance, 0.969; dPR, 0.967; frequency of equivalent or better values = 0.005: number 387/600: relation to SGC – 4 altered assignments, 14 unassigned, but preserves chemical order. 20 encoded functions.

**Figure 10C.**
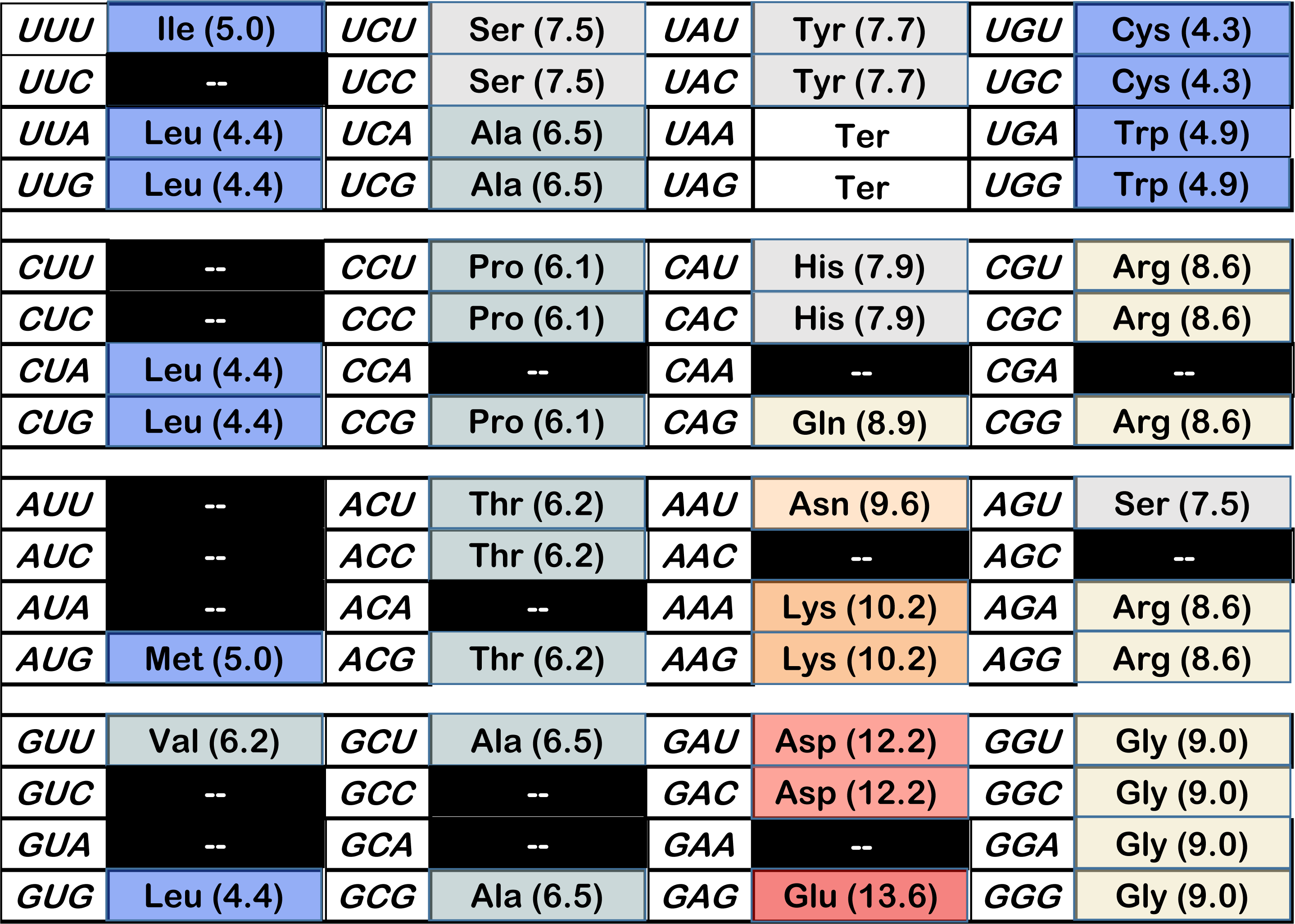
Example, continuous wobble joint progress ca. 0.95. spacing progress, 0.956, distance, 0.941, dPR, 0.965; frequency of equivalent or better values, 0.012: number 372/600: relation to SGC – 5 altered assignments, 17 unassigned, moderate perturbation of chemical order. 20 encoded functions.

**Figure 10D.**
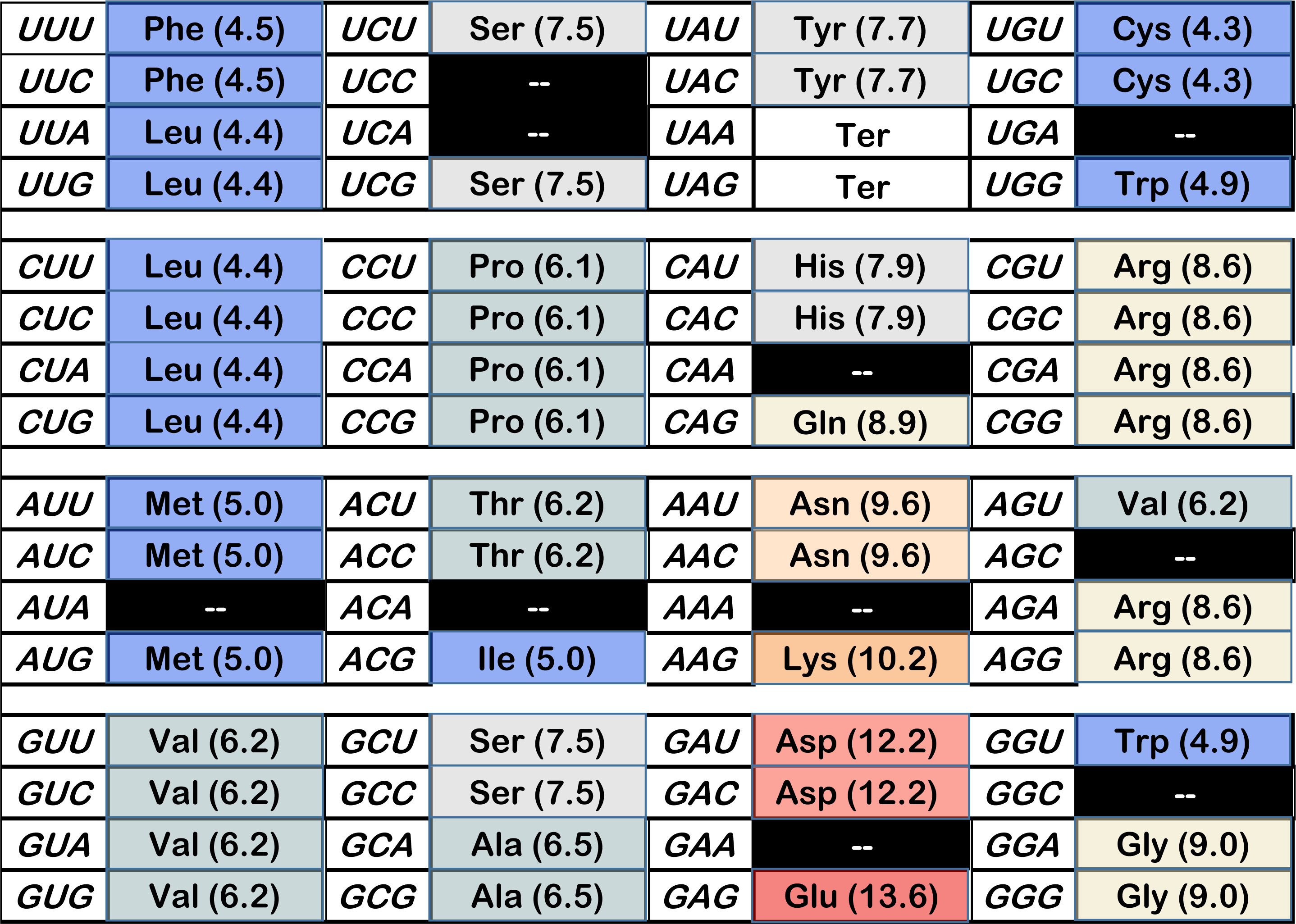
Example, continuous wobble joint progress ca. 0.9. spacing progress, 0.928, distance, 0.920, dPR, 0.909; frequency of equivalent or better values, 0.029: number 478/600: relation to SGC – 7 altered assignments, 10 unassigned, substantial perturbation of intermediate polar requirement bloc. 20 encoded functions.

**Figure 11.**
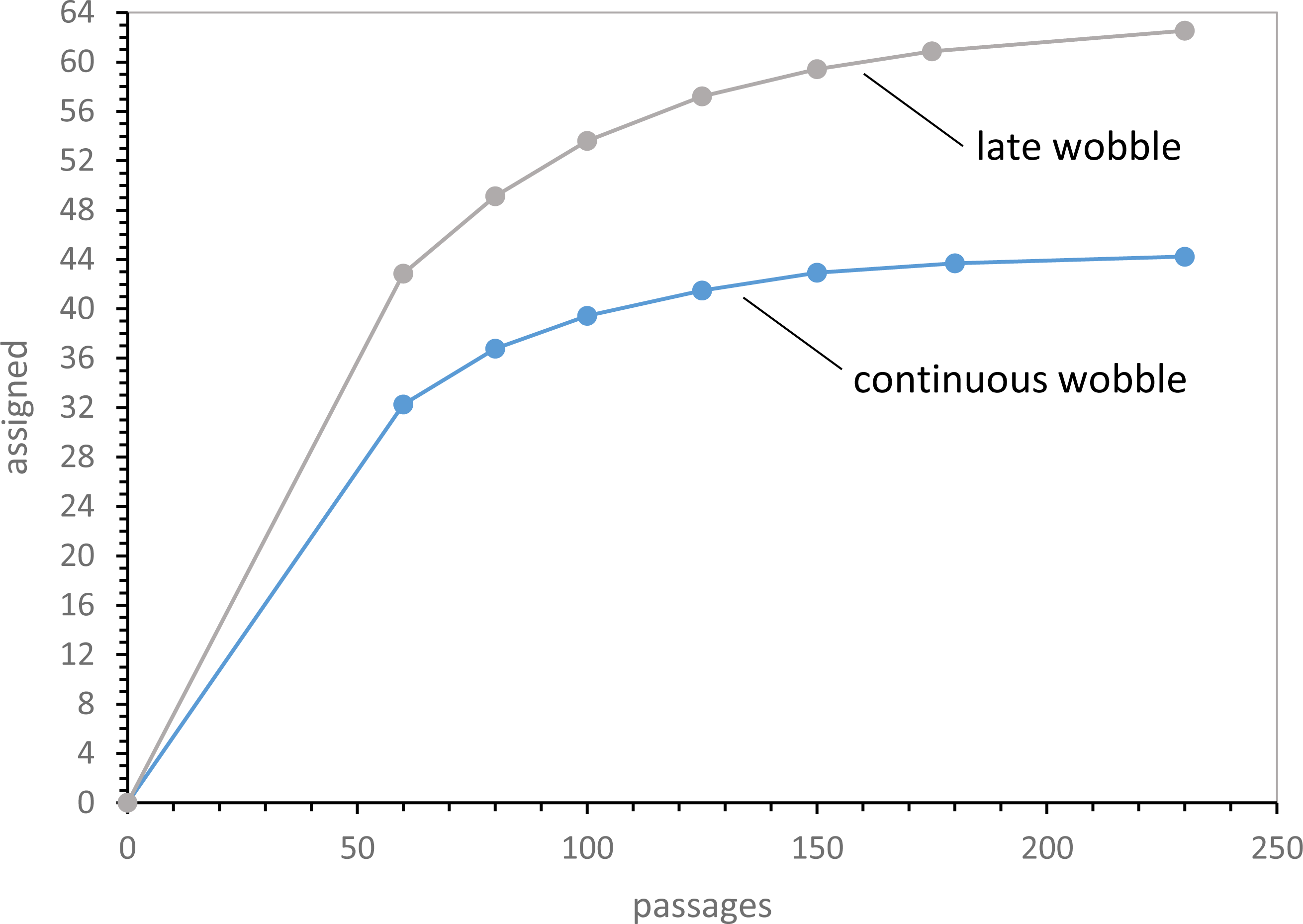
Number of codons assigned early during late and continuous wobble code evolution. 1000 evolutions with Coevo_PR assignments, Pmut = 0.04, Pdecay = 0.04, Pinit = 0.60, Pwob = 0.5, Prand = 0.1.

**Figure 12.**
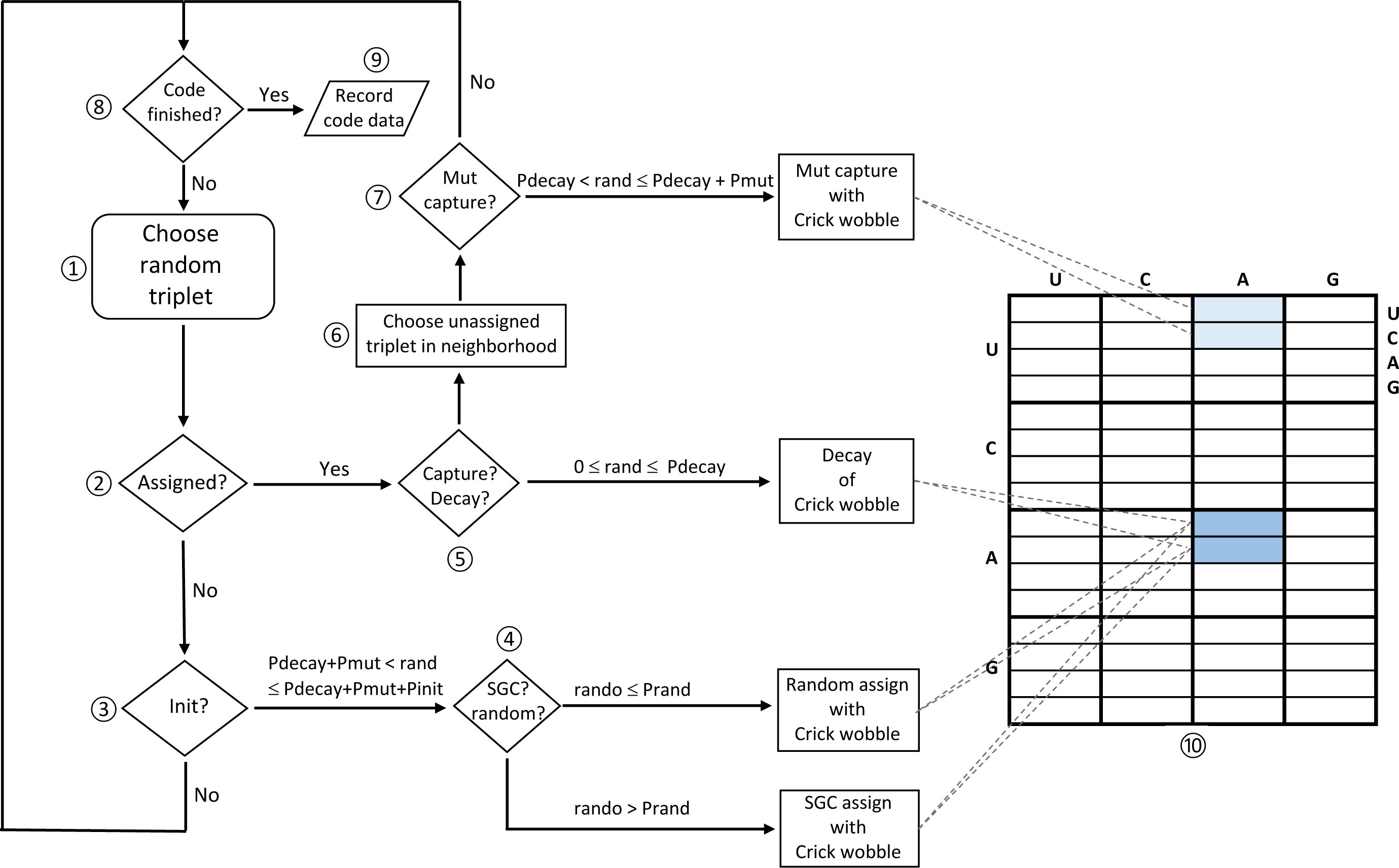
Operations during one computed passage. The logic of coding evolution during one computer passage through a nascent code is shown on the left, and the effect of these operations is visualized on the right as a typical coding table (as in Fig. 1, 9, 10) evolving with Crick wobble assignments (see “**Flow during one passage**..”, Methods).

These assumptions emphasize mode and kinetics during SGC approach, and de-emphasize mechanistic detail. Therefore this analysis foregoes some kinds of knowledge, to emphasize other kinds. I argue below that new resolution of the likely route to the SGC results, without requiring still-unknown mechanistic detail.

### Relations between identical and similar functions

Examination of triplets occupied by similar or identical amino acids in the SGC suggests regular relations between multiple assignments for similarly encoded functions.

### Third codon position

As has long been evident (Woese 1965), third codon positions often vary without changing coding, producing XY **A/G**, XY **U/C**, XY **U/C/A** or XY **U/C/A/G** blocs with similar assigned functions and polar requirements. This is not likely due to mutational uniqueness in third position triplet nucleotides, which presumably mutate as do other nucleotides. Instead similarity is attributable to wobble (Crick 1966), which assigns versatile base-pairing to third codon positions, reading them by ambiguous pairing with the same molecule. So, the code easily expands to accommodate third position mutational variation, immortalizing many such easy SGC expansions (Fig. 1A). SGC structure implies that that contemporary acceptors wobbled because wobble (Crick 1966) assignments are frequent.

Such order extends to amino acids that are not identical, but similar chemically, judged by polar requirement. Whenever a code box containing XY **U/C/A/G** also contains different amino acids, the amino acids have similar polar requirements but varied chemistry. This is true for chemically varied amino acids: hydrophobics like Phe and Leu, weakly polars like Ser and Arg, and very polar side chains like Asp and Glu (Fig. 1A).

### First position

Less frequently, mutational variation in the first position appears to have been captured: for example, with identical residues as for Leu UU **A/G** and CU **A/G**, or similarly, Arg CG **A/G** and AG **A/G**. Again, vertical columns of the same color (similar PR) often join otherwise chemically different amino acids by first position change. Gly-Arg, Tyr-His and Ser-Arg are examples (Fig. 1A).

### Second position

Least frequently, the SGC suggests capture of second-position variation for an identical function, the clearest possibility being UAA/UGA terminators. However, relations between chemically similar amino acids via second position change are common in the SGC, as for Ser-Tyr (Fig. 1A).

### The formative influence of mutational neighborhoods

These observations are consolidated by supposing that code evolution was guided by likely mutational pathways. A triplet with a given function might transfer function to a triplet related to it by single mutation. Thus there are three possible triplets that might be captured at the first, second and third triplet positions; nine possible captures in total. These nine changes comprise a triplet’s “mutational neighborhood”. When neighborhood mutations were readily accommodated, as at wobble positions, the code frequently expanded by that route.

Here, we simplify by assuming that all mutations are equally likely, though there is evidence that transitions (pyrimidine to pyrimidine and purine to purine) are more probable than transversions (purine to pyrimidine, or its reverse; Lehman and Joyce 1993; Vawter and Brown 1993; Collins and Jukes 1994; Kumar 1996).

### Plausible primordial acceptor RNAs from selection experiments

Selection-amplification for aa-RNA synthesis from its natural aa-adenylate precursor readily yields small aa-RNA producing catalysts (Illangasekare et al. 1995). By selecting aa-RNA synthesis without requiring aminoacylation of an arbitrary 3 ′ sequence, such an RNA active center can be reduced to a 5-nt ribozyme aminoacylating a 4-nt substrate RNA (Chumachenko et al. 2009) with only 3 nucleotides conserved for aminoacyl transfer. Thus the natural aminoacyl-RNA precursor, an activated amino acid adenylate, is bound and its amino acid regiospecifically esterifies the terminal 2 ′ hydroxyl of a tetramer RNA within a tiny RNA active center (Yarus 2011). The dimensions of such a catalytic RNA pentamer are not large enough to surround an amino acid, and indeed the small aminoacylator is not amino acid specific (Turk et al. 2011).

Varied selection data show that sidechain-specific amino acid binding RNAs exist, and require a minimum of 18-20 ribonucleotides (Yarus et al. 2005a; Yarus 2017b). Thus, regiospecific aminoacyl transfer requires a surprisingly simple center with only three conserved ribonucleotides. Ribonucleotides therefore are unexpectedly proficient at trans-aminoacylation catalysis. In pronounced contrast, many more nucleotides would usually be required to add side chain specificity. Therefore, amino acid specificity is not expected in the very earliest, small aminoacylation catalysts (but see (Illangasekare and Yarus 1999)).

Accordingly, selection-amplification suggests that the simplest, therefore earliest, ribozymic aminoacyl-RNA synthetase would catalyze RNA-specific acylation, via 3 or more specific base-pairs to an oligonucleotide acceptor, but would transfer multiple amino acids. The small aminoacyl-ribonucleotide product, using its free pairing nucleotides, could also base pair relatively specifically with a subset of codons (Illangasekare and Yarus 2012). The aminoacyl-RNA would thereby associate its triplet codon(s) with a set of amino acid sidechains. Base pairing nucleotides that bind RNA substrate to ribozyme can be changed with only small effects on activity (Illangasekare and Yarus 2012). So, mutation of a base-pairing, proto-anticodon nucleotide would allow the acceptor oligonucleotide to base pair with a new set of codons. New codon specificity therefore requires only a synthetase duplication and a single base pairing mutation. Such mutant aminoacyl-RNAs associate their amino acids with neighboring triplet(s), the event here termed mutational capture (see Methods, Fig. 12).

Further, ribonucleotides can be added to the small, unspecific aminoacylation active center. Extensions at both ribozyme and acceptor termini permit continued catalytic activity (Illangasekare and Yarus 2012; Xu et al. 2014). Such nucleotide additions might permit a new fold that allows amino acid sidechain specificity. For example, sidechain-proximal nucleotides potentially restrict large amino acids, making aminoacylation selective for small side chains. So, with two sequence changes (a proto-anticodon change and one proximal to the sidechain), previously untranslatable triplets might acquire a novel meaning, chemically related to a pre-existing assignment.

### Aminoacyl-RNA summary

Existing molecular data suggest a primitive manifold of specific acceptors, reading restricted codon groups, but using a single common aminoacyl transferase catalytic center, whose ribozyme can be as small as 5 ribonucleotides (Illangasekare and Yarus 2012). This RNA can be elaborated to add amino acid selectivity. Acquisition of new triplet specificities without losing aminoacylation activity permits such an aminoacyl transfer center to readily explore its triplet neighborhood, capturing the nine codons in its mutational neighborhood; that is, lying a single mutation away.

### Simple wobble

Early wobble coding must be minimal, independent of complex nucleotide modifications which can only arise later (Grosjean and Westhof 2016). To model wobble, I use a potentially primitive system (Crick 1966), requiring only natural nucleotides. In particular, third position G:U wobble pairs are allowed. Acceptor (anticodon): coding (codon) pairs include A:U, G:U, G:C, U:G, U:A, and C:G. Table I lists these and allows visualization of mutational transitions, and therefore of the evolutionary routes that simplified wobble coding most likely will follow. Thus, for example: one cannot assign XYA or XYC specifically; such functional triplets exist only as members of wobble pairs. If a wobble or non-wobble choice is made, as for codon XYU, wobble occurs with probability Pwob.

**Table I.** The simplified Crick wobble system. **Y** and **Z** are arbitrary nucleotides. Third acceptor complement position **G** may specify coding **C** or **U**; third acceptor complement position **U** may specify either coded **A** or **G** (Fig. 12). Coding triplets are permitted only one acceptor.

|  |  |  |  |  |  |  |  |
| --- | --- | --- | --- | --- | --- | --- | --- |
| Acceptor | 3' U |  | 3' C |  | 3' A |  | 3' G |
| Coding 1st | 5' AYZ |  | 5' GYZ |  | 5' UYZ |  | 5' CYZ |
| Acceptor | 3' U |  | 3' C |  | 3' A |  | 3' G |
| Coding 2nd | 5' YAZ |  | 5' YGZ |  | 5' YUZ |  | 5' YCZ |
| Acceptor | 3' U |  | 3' C |  | 3' A |  | 3' G |
| Coding 3rd | 5' YZA |  | 5' YZG |  | 5' YZU |  | 5' YZC |
| Coding 3rd | 5' YZG |  |  |  |  |  | 5' YZU |

### Quantitative detection of evolutionary progress

To compare evolving coding tables, objective measurement of differences like those between Fig. 1A and 1B is essential. With the SGC (Fig. 1A) and the above discussion in mind, code order is measured using three progress indices.

### Mean mutational spacing between identical assignments (spacing)

We are interested in grouping of identical functions because SGC coding occurs in compact groups (Fig. 1A). Progress toward this condensed goal is measured by counting mutations required to superpose triplets for identical functions (amino acids and start/stops). This distance (termed “**spacing**”) is ≤ 3 mutations for every triplet comparison; = 3 if all three coding nucleotides must be changed. Further, each pair of triplets must be counted only once, not duplicated by starting from both participants. In practice, it is useful to normalize distances for the number of pairs, calculating mean distance/triplet pair. Normalization makes spacing resilient when tables with varying numbers of unassigned triplets are compared. In 1000 random complete coding tables, identical functions are 2.284 (mean) ± 0.002 (sem) mutations apart. The SGC has a mean distance of 1.30 mutations between identical functions by the same criterion. Thus spacing progress captures the SGC’s exceptional compaction – tracking progress from random tables (spacing 2.284) toward the condensed SGC (spacing 1.30).

### Distance from the SGC (distance)

Progress to any code is of interest, but most particularly, progress toward the SGC. Distance to the SGC is quantified by totaling the total number of mutations required to move from triplets in a novel table to triplets for identical functions in the SGC. Again, only identical functions are compared, all possible pairs are counted once, and the result is normalized to yield the mean distance per triplet comparison. One thousand independent completely random tables average 2.286 ± 0.002 mutations from the SGC (per pair) by this measure. Random spacing and distance are further clarified in Methods.

### Chemical order (dPR)

The SGC shows exquisite, virtually comprehensive ordering of amino acids by polar requirement (colored areas, Fig. 1A). We quantitate chemical order by summing absolute polar requirement differences over all amino acid pairs in mutational neighborhoods, using corrected amino acid polar requirements (Mathew and Luthey-Schulten 2008) closely related to those measured chromatographically by Woese (Woese et al. 1966). Only neighborhood pairs that differ are counted and normalized for the number of comparisons. Thus, dPR does not overlap with spacing: dPR counts only non-identical residues. So, dPR specifically measures chemical grouping, not coding proximity. This normalized distance is 2.98 ± 0.01 per amino acid pair (in polar requirement units) for 1000 random tables versus 2.069 for the SGC, thereby allowing dPR to report chemical order (Fig. 1A). dPR is the only progress index that explicitly utilizes the notion of mutational neighborhood.

### Indices of coding order: progress values

In order to make progress indices transparent indicators of SGC proximity, they are used in a form which does not require comparison to other numbers. This “progress value”, is 0.0 for unordered, random coding tables and 1.0 when order equivalent to the SGC is attained. Thus progress from random coding to SGC order appears as decimal zero to one respectively; progress value is the fraction of mutational or chemical distance to the SGC covered.

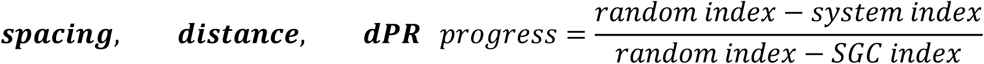

Progress values can be < 0 or >1 because systems can be more disperse than random coding tables or more frequently, more ordered than the SGC itself. Still, mean decimal spacing, distance and dPR allow assessment of a calculation yielding tens of thousands of numbers, indicating whether greater or lesser mean SGC likeness was attained. Thousands of such discriminations were made in the present inquiry. While progress might be differently defined, below these definitions do locate the SGC.

### Progress values respond to random assignment

To clarify progress values, Fig. 2A plots mean spacing, distance and dPR for groups of 250 full coding tables constructed with varied numbers of randomly-chosen SGC assignments. Unassigned triplets were filled with random assignments with no set relation to the SGC. These coding table populations therefore are otherwise random, but have a specified fraction of SGC ordered assignments - the latter fraction is plotted across Fig. 2’s x-axis. Distance progress is accurately proportional to the fraction of random triplet assignments, starting from the SGC at upper left. dPR and spacing progress are more sensitive to random assignment, declining to near-random values before all triplets are randomized. Spacing is most sensitive to random triplet intercalations, but all three progress indices respond progressively to small deviations from the SGC, rationalizing their use to assess SGC proximity.

### Progress values have varied relations to SGC order

In Fig. 2B is shown spacing, distance and dPR progress for 10000 coding tables evolved by late wobble, a code history we will later dissect in detail. All distributions are roughly symmetrical single peaks and so well described by means and standard errors used here. Almost all evolved coding tables have progress value distributions highly shifted from random assignment (0.0), with means just below the SGC (1.0), indicating overall effectiveness for a late-wobbling evolution.

Full distributions in Fig. 2B also show that distance is the sharpest peak and finds the fewest codes at or exceeding SGC behavior. Spacing is broader and has an intermediate-size foot at or beyond the level of SGC grouping of identical functions. The broadest and least symmetrical distribution is for dPR and chemical order among closely related triplets. As one result, chemical order is often the most frequently evolved in this population.

### SGC-like codes require three upper tail properities

Moreover, Fig. 2B shows that access to realistic coding tables is very sensitive to underlying coding order. Tables resembling the SGC are those simultaneously in the upper tails of three progress distributions. Fraction of evolved tables with three progress values near 1, the fraction with SGC order, will therefore vary rapidly and non-linearly when change in history shifts or spreads underlying spacing, distance and dPR distributions (Fig. 2B), even slightly

Thus, the joint distribution of progress values (Fig. 2C) quantitatively implements the present biological goal; a code that possesses SGC-like grouped assignments, SGC-related and chemically ordered (Fig. 1A).

In Fig. 2C the fraction of coding tables with joint progress greater than the abscissa value is calculated. For example, ≈ 50% of evolved coding tables have spacing, distance and dPR simultaneously >= 0.7. More specifically, when the “vicinity-of-the-SGC” is desired, the region under the rightward gray bar will be utilized. In other words, those coding tables with spacing, distance and chemical order simultaneously covering >= 90% of the distance from random codes to the SGC will be cited, to target discussion. Solid and dashed curves in Fig. 2C represent two ways to evolve a genetic code. Below, we will return to these data.

### The code evolution model repeatedly completes codes

A biologist may be slightly interested in averaged behavior for coding table populations. Such an aggregate accurately follows underlying kinetic rules, but for example, never finishes a coding table, persisting forever in an average near-steady-state with unassigned triplets. In contrast, the coding table subset that evolves to finish a code is of immediate interest. Completely finished codes assign all 64 triplets (termed “full” tables) or encode all 22 functions (termed “complete” tables).

Coding history is computed (described in Methods, Fig. 12) by following one coding table at a time to its particular fate. A triplet is randomly chosen. Subsequent events occur at random on the basis of probabilities for initial triplet assignment (Pinit), mutational capture of a nearby triplet by existing assignments in an assigned triplet (Pmut) or assignment decay (Pdecay). Using randomized numbers to choose chance events with specified probabilities, a chosen unassigned triplet can be allocated to one of 22 essential functions (Pinit). If the randomly chosen triplet has already been assigned, it can capture new triplets for its function, or related functions, via neighborhood mutations (Pmut). Alternatively, its function can decay, with the triplet losing its previously assigned meaning, to become unassigned again (Pdecay). Probabilities are chosen to limit outcomes to a total probability of ≤ 1.0. Repeating such chance events, only one of which occurs in a passage, ultimately builds a code with desired properties: for example, full (64-assignment) codes, or a complete (22 function) code. Repetition of such computations compares evolutionary routes by determining the probability of an SGC-like outcome.

### Kinetics by choosing random triplets

Because it is not usual to compare rates by performing a succession of random events, we first show that this procedure yields expected kinetic behaviors. Fig. 3 exhibits velocities for initial triplet assignments, mutational captures and assignment decays as a function of the number of assigned triplets (randomly chosen from the SGC in 1000 repetitions). First order reactions should be proportional to reactant availability, second order reactions to the product of two reactant availabilities.

### Initiation

Initial assignment linearly declines in rate as triplets are filled with random SGC assignments. Initiation is a maximum with no triplets filled (at left, where the least squares line extrapolates to the complete table’s Pinit = 0.6), decreases linearly as triplets become assigned, and extrapolates near zero when 64 triplets are occupied, so that no initiation can exist. Thus: accurate first order initiation is seen.

### Decay

Assignment decay should also be first order in assigned triplets. It is zero at left (where there are no assignments to decay), increases linearly as assigned triplets increase, and extrapolates to a maximum reflecting the probability of table decay itself (0.04/passage) when 64 triplets are occupied, and therefore full probability of decay is expected. Thus: accurate first order decay is observed.

### Mutational capture

Transfer of an assignment to a neighborhood triplet contrasts with initiation and decay: it requires an assigned triplet and an unassigned one, the latter to be captured for a similar assignment. Expansion of the code by mutational capture therefore should be second order, varying with the product (assigned*unassigned) triplets. The data shows that the expected second order maximum capture rate is observed when half, 32 triplets, are assigned. Moreover, the fitted rate extrapolates to zero both when assigned codons = 0 (at left) and also when unassigned codons = 0 (at right). So, mutational triplet capture behaves as a second order reaction. There is further quantitative support for this rate analysis in Methods.

One might summarize Fig. 3 by saying that computer passages through an evolving coding table are proportional to time. But there is nevertheless a difference in definition. To respect this difference, durations are expressed in “passages” (computational transits of a nascent coding table) rather than “times”.

### Two eras in a nascent coding table

By putting off some details (of mutational capture mechanisms), coding table fates can now be computed. Figure 4A shows mean data for a population of 1000 coding tables without wobble (characteristics in legend), through their initial 4096 passages. There is an initial period of rapid change (≈ 0-200 passages), then a near steady-state in which assigned and unassigned triplets change little. In that later era, total decays, initial assignments and mutational captures increase at almost constant rates, but mean assigned/unassigned triplets are almost constant. Assignments are rapid initially (because many unassigned triplets exist), decays increase after a delay (in which assigned triplets accumulate), and mutational captures accelerate, then slow as requisite unassigned triplets become rare. Ultimately assignment events (initiation and mutational capture) and decay events balance, and a steady state emerges. In Discussion we return to persistent unassigned triplets, and to stably incomplete coding, exemplified in Fig. 4A as a mean of 60 steadily assigned and 4 unassigned codons.

Fig. 4B, however, shows new, finished code behavior. Both transient and near-steady-state behavior appear also for full and complete coding tables of biological interest. Full coding tables (all triplets assigned) appear after a delay and then are stable at about 0.016 of the population. Complete coding (22 encoded functions) both appears earlier and also is more abundant: ≈ 0.22 of all tables. This sequence reflects the fact that ≥ 22 events minimally complete a table, but ≥ 64 events are required to fill a table. Two distinct eras: early transient emergence and later stable codes, shape code evolution and accordingly, figure again below.

### Sources for steady state order

Now consider evolution of progress values. Fig. 4B shows spacing, distance, and dPR progress with increasing Fig. 4A duration. Coding progress also has a steady state. Progress values are similar at all points in a population’s history, once coding tables are substantially occupied. This is equally true for spacing, distance and dPR. Thus progress is near-constant in time for tables with the same transformation probabilities (Fig. 4A, 4B). Finally, Fig. 4 evolution history employs random initiations and random later mutational captures. Fig. 4B extensively documents the lasting random fate of such a table (progress values ≈ 0) throughout 4096 passages.

Thus, to evolve an SGC we must include source(s) of order. Order comes from assignment of early triplets matching the SGC (as for stereochemical origins). Such initial stereochemistry will be insufficient, but its failure defines more successful code evolution.

### Coding tables with initial SGC assignments

Consider coding that begins with 16 randomly-chosen SGC triplets. Using different sets of initial SGC triplets averages effects of particular dispositions. Fig. 5 presents such average passages to varied levels of encoding, including completion at 22 encoded functions. The number of initial triplet assignments required to attain final levels of encoding (in addition to the initial 16) is also shown.

Because results differ greatly in wobbling (Fig. 5A, 5B) and non-wobbling (Fig. 5C, 5D) coding systems, for the first time Fig. 5 also distinguishes coding systems. Non-wobble codes, like those treated thus far, admit any assignment to triplets, however their codon sequences may be related. In contrast, wobble (Table I) allows G:U and A:U third position pairs (Crick 1966), fixing some adjacent codons’ meaning.

### Coding tables initiated with 16 SGC assignments: wobble coding

Fig. 5A shows mean durations (passages) and number of random assignments (inits) needed to attain particular numbers of encoded functions. Notably, both time and assignments needed to reach a specified wobble code complexity increase dramatically after 20 encoded functions. In fact, starting at an initial mean of 12.35 functions (from 16 chance SGC triplets), encoding the last two functions costs more than 100-fold as much time and assignments as do the other 20 encodings. This implies a history of great complexity - a complete wobble coding table has assigned triplets an average of ≈ 25000 times, thereby overwhelmingly making repeated, futile assignments.

This is ‘completion complexity’, a reflection of the difficulty of fitting together wobbling coding boxes in a fixed space that must contain 22 of them. Many explorations, involving decay and reassignment (detected as inits in Fig. 5A), are required to complete a wobble coding table. In addition, forces driving change weaken as full tables are approached. Initiation slows near completion because unassigned codons become rare (Fig. 3). Mutational capture also slows near completion because one participant, the unassigned triplet, also becomes rare (Fig. 3). Finally, decay of assignments will be maximal near completion, also opposing completion (Fig. 3).

Moreover, there are accompanying effects on wobble coding order. As random assignments are added, progress decays (Fig. 5B) and coding tables move away from SGC order. However, there exists a partial exception; spacing. Because wobble initiations make closely-spaced identical assignments, wobble’s spacing progress uniquely resists erosion, persisting indefinitely at ≈ 40% of SGC levels. However, wobble’s spacing order occurs without parallel effects on distance or dPR, which descend to indistinguishability from random coding (line labeled ‘random’).

But note the contrasting situation *before* any evolution. Random groups of 16 initial SGC-like assignments (at ‘init’) have average progress values (points at upper left) that approximate the SGC itself (line labeled SGC). Initiation with wobble particularly improves spacing and dPR order (compare Fig. 5B, 5D). Sixteen such wobble initiations immediately make an incomplete coding table with spacing, distance, and dPR similar to the SGC.

### Coding tables initiated with 16 SGC assignments: non-wobble coding

Fig. 5C shows mean duration and number of random non-wobbling assignments to reach particular numbers of encoded functions. Completion complexity exists for non-wobble codes, but is much less obstructive than with wobble. To pass from 20 to 22 non-wobbling functions (Fig. 4C), only 2.9-fold more initiations and 9.7-fold more duration is required. In addition, even if non-wobbling completion at 22 functions is mandated, only ≈ 1 additional assignment per triplet must occur.

However, code completion again undermines progress values and coding order. Fig. 5D shows that spacing progress is generally reduced because the close spacing enforced by wobble assignments does not exist (compare spacing lines, Fig. 5D and 5B). Keeping in mind that non-wobble evolution is far shorter (Fig. 5C vs 5A), requiring 1/1000 the 20 -> 22 function assignments for wobble - spacing, distance and chemical order still descend to near that of randomly-formed coding tables (Fig. 5D).

### Wobble summary

Non-wobble completion is quicker and simpler than for a wobbling code, but if supplied by initial SGC assignments, code order still decays decisively. The evolutionary history modeled in Fig. 5 (initial stereochemistry, with and without wobble) is improbable, even if one’s goal is coding that only faintly resembles the SGC. One must avoid the decay of SGC-like order supplied by initial wobble assignments (Fig. 5B, 5D), and also mitigate related effects of delay (Fig. 5A, 5C) during progress from a near-complete to a complete wobble code.

### An apparent solution for completion complexity

Dramatic delays are confined to the era between 20 encoded functions and completion (Fig. 5A). Accordingly, it is possible that a minority of encoded functions evolved later than the majority, perhaps via a different route. This is an appealing notion for independent reasons.

Coding of translational initiation differs greatly in bacteria and eukaryotes (Kozak 1999). Bacteria initiate internally, using mRNA-rRNA complementarity as a guide, while eukaryotes scan from a 5 ′ mRNA end to a first favorable AUG (Hinnebusch and Lorsch 2012). These fundamental differences suggest that translation initiation evolved late, after divergence of the major domains of life.

Translation termination also differs in bacteria and eukaryotes, much more than encoding of amino acids, which is similar throughout Earth biota. Protein release factors have different evolutionary origins in different domains of life (Vestergaard et al. 2001), and auxiliary factors, like those that recycle the joined ribosomal subunits after termination, are also of independent evolutionary origin (Zavialov et al. 2005). Moreover, definitive termination factors are sophisticated protein catalysts (e.g., Adio et al. 2018) that cannot exist until translation itself is sophisticated. Such considerations suggest that translation termination also took its final form late, after separation of life’s domains (Burroughs and Aravind 2019). Thus the suggestion of a majority of quickly encoded functions (≈ 20) and a small number added later by a different logic (≈ 2) has extensive, long-standing molecular support.

### Rapidly evolved codes with wobble

Fig. 6A shows that average coding behavior (as in Fig. 5) conceals a possible resolution for wobble’s completion complexity. Fig. 6A plots the distribution of times to acquire 20 coded functions, for wobbling and non-wobbling codes, in successive 50-passage windows. Firstly, evolution to 20 functions (Fig. 6A) makes wobble less burdensome: mean times (signpost-shapes) to code completion are 28 fold greater for 20-function wobble codes than without wobble, instead of 1000-fold (Fig. 5) at 22 encoded functions. Modes, most probable completion times, do not actually differ greatly for wobble and non-wobble codes encoding 20 functions. Instead, wobble requires longer mean evolutionary times because of a long tail of tortuous histories, in which the many assignment decays and re-initiations (Fig. 5A) mentioned above in ‘…: **simple wobble coding**’ gradually occur. So: if most probable routes (left hand peak in Fig. 6A) are taken instead of average ones, codes that exhibit SGC wobble, but also appear quickly can evolve. Fig. 6B reinforces this discussion, showing that complete coding tables do not possess substantial early completions. A 22-function coding goal makes rapidly completed coding tables rare (Fig. 6B), instead of common (Fig. 6A).

### Late wobble

So, two short paths to wobble coding appear. In the first, 20 functions are encoded without wobble, exploiting the easy access non-wobble coding has to nearly complete tables (Fig. 5C). Modern ribosomes use complex rRNA conformational changes to limit codon: anticodon complexes to third-position wobble (Moazed and Noller 1990; Ogle et al. 2001). Moreover, the tRNA anticodon loop-and-stem structure has a complex role in translational efficiency (Yarus 1982), including suppression of errors (Ledoux et al. 2009). Thus, it is plausible that simpler base pairing was primordial, and that evolutionary advances in both ribosomes and adaptor RNAs were required to make accurate third-position wobble (Table I) possible. Late wobble innovation is quickly adopted - pre-existing 20-function codes quickly add wobble wherever possible. These events will be called “late wobble”.

### Continuous wobble

A second rapid route to SGC-like wobble coding, called “continuous wobble”, allows wobble assignments (Table I) at initiation of coding and throughout. This path seeks access to the SGC via the early subset of 20-function wobble codes (Fig. 6A). An SGC evolved via this minority is readily accessible - about ¼ of all code evolutions are elligible (Fig. 6A). However, SGC evolution via a subpopulation, even via a substantial minority like 25%, has consequences that return in Discussion.

### A second barrier: coding table order

Decline of progress values in Fig. 5B and 5D imply that the exquisitely ordered SGC (Fig. 1A) will require specific, persistent organizing influences. Therefore, we now compare often-cited sources of order. Calculations below compare 6 ordering mechanisms utilizing coevolution and paralogous selection, adaptation and neutral mechanisms. These 6 mechanisms (termed Coevo, Coevo_PR, 0±1 PR, 0±2 PR, 0±3 PR, 0±4 PR) shape capture specificities for new triplets acquiring assignments related to existing ones.

### Sources for code order

SGC non-randomness (Fig. 1A) is frequently attributed to stereochemical and/or historical causes (Knight et al. 1999).

### Stereochemistry

Stereochemistry implies that the amino acids and cognate coding triplets are related by chemical interaction (Woese 1967; Crick 1968). Thus, stereochemical hypotheses predict that contemporary experiments can reveal code origins by studying interactions. An example: RNA binding sites selected for amino acids contain cognate coding triplets with unusual frequency (Yarus 2017b).

### Coevolution

In another logic, historical explanations of coding order take one of three somewhat parallel forms. The first is **co-evolution**: the idea that ancient encoded amino acids ceded their codons successively to related amino acids produced via extension of biosynthetic pathways (Wong 1975). Co-evolution of the code and biosynthesis can be examined by testing the SGC to see if SGC triplet assignments are frequently connected in the way predicted by synthetic pathways (Amirnovin 1997; Ronneberg et al. 2000). Moreover, a possible molecular remnant of co-evolution exists (Di Giulio 2002).

### Adaptation

The second category of historical ideas is that there is a selective **adaptation** behind the code’s order. For example, minimizing polar requirement change might guide capture assignments by minimizing the structural effects of substitution errors on protein structure (Freeland and Hurst 1998a). The SGC is very effective in minimizing the cost of such errors (Freeland and Hurst 1998b).

### Neutral change: paralogy

A third **neutral** mechanism has been tested (Massey 2008, 2016, 2019). Because successor RNA-amino acid interactions would likely be molecular derivatives of prior RNA-amino acid interactions, they would employ related sequences. As a result, there could be sufficient order in a descendant coding table to explain the relatedness of triplets and SGC amino acids. Descent of related RNA sequences for related amino acids also occurs within adaptation. Selection therefore produces code order by means paralleling the neutral mechanism. Because adaptation and neutral paralogy plausibly exist together, producing overlapping, similar code order via a shared mechanism, I suggest their unification as paralogical sources of related triplet-amino acid assignments. Unification of selection and relatedness implements Crick’s prescient comment that “similar amino acids would tend to have similar codons” (Crick 1968).

### Encoding order: Coevolution

The above considerations, determining functions assigned to neighborhood triplets captured by an existing assignment, have been programmed. For coevolution, related triplets are assigned to amino acids linked by synthetic pathways, as suggested by Wong (Wong 1975), but using later thermodynamic corrections (Ronneberg et al. 2000). Such assignments for the purpose of testing co-evolution usually are restricted to unique biosynthetic pathways, and common amino acid interconversions are ignored. However, in present evolutions, common amino acid interconversions are included and used to guide assignment of triplets to related amino acids. Coevolutionary amino acid conversions used are listed in (Ronneberg et al. 2000). This assignment mechanism is called Coevo.

### Coevolution respecting PR chemical similarity

Here biosynthetically related triplet/amino acid assignments are made as for coevolution, but synthetically related amino acid assignments that best conserve polar requirement are chosen with higher probability, rising as PR difference decreases. This mechanism is called Coevo_PR.

### Selection and paralogy

To represent paralogical sources of order, related triplets are assigned amino acids with related polar requirements. Amino acids are ordered by their PRs (Mathew and Luthey-Schulten 2008), and related triplets are randomly assigned to the next amino acid, up or down, in the PR list (0±1 PR). Alternatively, random assignments are made ± 1 or 2 places in the PR list (0±2 PR), or randomly, ± 1, 2 or 3 places in the PR list (0±3 PR). When such random changes fall outside the range of real amino acid PRs, unoccupied triplet assignment defaults to the same amino acid as for the already assigned triplet (0 PR changes). These chemically conservative paralogical mechanisms are called 0±1 PR, 0±2 PR, 0±3 PR and 0±4 PR.

### Revised code evolution: interspersed SGC assignments

I now react to Fig. 5A-5D by making several mechanistic alterations, and by targeting 20-function codes to minimize completion complication. Ordered assignments implementing the SGC (as for stereochemical assignments) are not made solely at initiation of code history, but arbitrarily interspersed with random assignments, throughout code evolution. In this way, ordered code exposure to random dilution (Fig. 5B, 5D) is shortened. The probability of random initiation is Prand. (1 – Prand) is the probability of SGC-like initiation; both are constant throughout code history.

### First route: evolution to 20 functions, then late wobble

Time and assignments for late wobble are similar for all histories, requiring ≈ 50 initiations in ≈ 170 passages. Notably, quick, advantageous evolution is retained: coding tables with 20 functions appear 20 to 30 times faster than average for continuous wobble. Moreover, different assignment mechanisms under late wobble require few, and similar, initiations (≈ 0.78 assignments/ triplet). Thus, all late-wobbling assignment histories yield coding tables rapidly, without multiple decays and assignments. In fact, the 20 to 30-fold shorter times to late-wobbling coding tables are accompanied by similar-fold decreases in other events, like assignments transferred to new triplets. Equivalent time and inits among assignment mechanisms supports evolutionary choice by criteria other than rate or complexity.

### Assignment mechanisms and approach to the SGC

We now compare mean code order after varied assignment mechanisms during acquisition of 20 encoded functions. Fig. 7A shows average progress for six assignment histories.

The outcomes of different assignment mechanisms faithfully reflect their individual rationales. Paralogical modes 0 ± 1 PR, 0 ± 2 PR, 0 ± 3 PR, 0 ± 4 PR are defined to conserve polar requirement, and their net evolutionary effects reflect this definition. They indeed conserve chemical order better than coevolutionary modes, Coevo and Coevo_PR. As chemical conservation relaxes, 0 ± 1 PR to 0 ± 4 PR, chemically ordered final codes become less frequent. Chemical order (dPR) is always more attainable than grouping (spacing progress), with resemblance to the SGC (distance) always the least frequent. But if chemical order were the sole coding goal, paralogous assignments produce it most effectively (Fig. 7A).

But, notably: what assignment mechanisms neglect also matters critically. Conserving chemical order (dPR) alone ignores and therefore sacrifices spacing and distance. Coevolution within Coevo and Coevo_PR always yields more compact spacing and closer approach to the SGC. The most balanced choice is Coevo_PR (Fig. 7B). Its dual emphasis on both biosynthetically related assignments and related chemistry, yields the most frequent mutual access to SGC-like spacing, distance and dPR together, though chemical order is still most easily attained. To facilitate this balance, Coevo_PR is employed in further examples.

We confirm this choice and also take a step toward realism, plotting a quantity more relevant to code evolution than average progress – the fraction of evolved tables with joint progress >= 0.9 (Fig. 2C). In Fig. 7B, abundance of coding tables in the SGC vicinity is similar for all paralogical modes: the differences in Fig. 7A are compensated by changes in distributions. However, coevolutionary assignments produce ≈ 2-fold more SGC-proximal codes, with Coevo_PR again the best.

### Second route: evolution to 20 functions with continuous wobble

Assignment modes Coevo, Coevo_PR, 0 ± 1 PR… act similarly in continuous and late wobble. Again, paralogical modes promote chemical order (Fig. 7C), with tight conservation (0 +/- 1 PR) more effective than less constrained assignments (0 +/- 4 PR). But spacing and distance order are again enhanced in coevolutionary modes, with Coevo_PR again benefitting from its built-in preference for chemical order.

And again (Fig. 7D), using joint progress as a more comprehensive indicator, paralogical modes are similar and more than 2-fold less productive of codes with SGC-like order than coevolutionary assignments. In Discussion, we consider the ≈ 50% superiority of coevolution with continuous wobble (Fig. 7D) over coevolution with late wobble (Fig. 7B).

### Near-complete codes from late wobble: random assignments

Disruption by random assignments first appeared in the specially constructed coding tables of Fig. 2. In Fig. 8A, mean disruptive effects of random assignment on 20 function late wobble code evolution are plotted. All progress is decreased by random assignment (upper curves, Fig. 8A). But as expected (Fig. 2B, 2C), disruption is greater when SGC-likeness is assayed using joint progress (Fig. 8A, lower curves, rightward ordinate). In particular, balanced progress gained from Coevo_PR assignments (Fig. 7A, 7B) is disrupted by a minority of random assignments. Good joint progress at Prand = 0 (no random assignment) is lost if more than 15% of assignments are random rather than identical to the SGC. While some random assignment is allowable, > 15% is incompatible with an SGC-like result, particularly for spacing and distance order. Thus, 10% random assignment was chosen for illustrative calculations above.

### Near-complete codes from continuous wobble: random assignments

Fig. 8B plots the effect of random assignment on continuously wobbling codes. In particular, it reinforces the previous limit: random substitution with continuous wobble must be restricted; certainly <= 15%, better <= 10%.

But Fig. 8B differs from the late wobbling case in Fig. 8A, as seen in its lower joint progress curves. Code order is similarly sensitive to random codon assignments in late and continuous wobble (joint progress >= 0.9; Fig. 8A, 8B). However, if coding capacity for >= 20 encoded functions is also required (joint progress >= 0.9 & >= 20 functions), then late wobble supplies more candidates than continuous wobble. This result traces to Fig. 6A: because random assignment’s effect on order is similar for late and continuous wobble, it is limitation to a minority (Fig. 6A) of continuous wobble coding tables that diminishes the frequency of codes in the vicinity of the SGC. By comparison, codes encoding 20 functions before adopting wobble are already near-complete: thus joint progress, and joint progress with completeness, superpose for late wobble (Fig. 8A).

### Evolved examples distributed coding outcomes

Because evolutionary outcomes span a large stochastic range (Fig. 2C), we must now grapple more fully with variation. To illustrate how progress statistics represent the SGC, coding tables under the rightward gray bar of Fig. 2C are displayed in Fig. 9. These examples were picked from 600 successively evolved random tables. The best available is shown, and also progress around 1.0, 0.95 and 0.9 to illustrate the nature of differing joint distributions.

Fig. 9 shows coding tables in descending order of joint progress: using SGC-like initiations, random assignment 10%, late wobble, and Coevo_PR controlling related triplet assignments. The most ordered table cannot be accurately placed because there are no comparable tables to define its real frequency; but frequencies for 1.0, 0.95 and 0.9 examples can be computed from their positions in the observed joint distribution (Fig. 9).

These tables exemplify the use of progress indices to characterize less-than-SGC order. For example, comparison to the SGC shows that tables 9A through and including 9D resemble the highly ordered SGC (Fig. 1A) much more than they do a random coding table (Fig. 1B), thereby substantiating progress index shifts plotted in Fig. 8A and 8B. A detailed examination of these examples also indicates that a code with high resemblance to the SGC would be accessible from a small population of hundreds of codes evolved by these means. Call this outcome ‘distribution fitness’, to indicate that the better members of a distribution contribute disproportionately to evolutionary potential. For example, about 1 in 24 late wobbling, 10% random, Coevo_PR coding tables is equivalent or better than Fig. 9D, which shows progress values ≈ 90% the distance from random to SGC coding.

Other non-trivial implications appear from Fig. 9. The frequency of coding tables with spacing ≅ distance ≅ dPR ≥ 1 is low (Fig. 2C, 7D). Thus, orderly coding tables are not a subset having uniformly favorable properties; instead, progress values vary individualistically.

### The second route to an SGC: continuous wobble to 20 functions

The second route to an ordered wobbling SGC is code completion during the early 20-function peak (Fig. 6A). In order to present explicit quantitation, we concentrate on a population of coding tables at 200 passages. Because all such coding tables have existed for exactly 200 passages, all experience similar mean development, with close to 51 initiations, 0.45 decays and 12 mutational captures.

### Overall order is roughly similar

As Fig. 2C showed, distributed joint progress for continuous wobble early is very similar to joint progress for late wobble, but with a slight advantage to continuous wobble. This continuous wobble advantage is assessed in Discussion.

### Assignment effects for continuous wobble

Order due to various kinds of mutational capture (Fig. 7A, 7B) also varies similarly to that for late wobble (Fig. 7C, 7D). Paralogous mechanisms conserve chemical order best, with tighter paralogous constraints (e.g., 0 +/- 1 PR, 0 +/- 2 PR) more effective. Again, coevolutionary mechanisms, Coevo and Coevo_PR, are better balanced, with better distance and spacing, and good, but usually less effective, chemical ordering. Thus we continue using Coevo_PR for specific continuous wobble calculations.

### Sensitivity to random assignment

The sensitivity of continuous wobbling to random (rather than SGC) assignments is again pronounced (Fig. 8B). All three progress values decline, with joint progress approaching random codes at Prand > ≈ 0.15, thus resembling late wobble (Fig. 8A). Moreover, joint overall order for continuous wobble is particularly sensitive to random assignments (Fig. 8B), as it was for late wobble (Fig. 8A), because of a similar requirement for simultaneous upper-tail behavior in three distributions. Thus, majority SGC-like initiations (Fig. 2, Fig. 8A) are not unique to late wobble, but similarly required for continuous wobble (Fig. 8B).

### A highly significant difference between continuous and late wobble

But late and continuous wobble coding differ. This is apparent in examples from 600 continuously wobbling (Fig. 10) versus parallel late wobbling (Fig. 9) coding tables. Fig. 10 illustrates the best order observed in a continuous wobbling population of 600 (joint progress values >= 1), and also codes with joint progress ≈ 1, 0.95 and 0.9.

More frequent unassigned triplets (black with white dashes) among continuous wobbling tables (Fig. 10) are apparent, compared to late wobbling (Fig. 9). Example tables were chosen to illustrate joint progress, but excess unassigned codons are not due to human choice. Fig. 11 shows that a ≈ 20-triplet assignment superiority for late wobble is not idiosyncratic and will not disappear; it is characteristic of the near steady-state. Evolution of almost-complete 20 function wobble codes will leave about a third of amino acid triplets unassigned. In contrast, unassigned late wobble triplets are fewer, more comparable to the small number of yet-to-be-encoded functions.

## Discussion

### The major conclusion

A computation is introduced to evolve finished coding tables. Evolutionary qualities are varied to evaluate coding pathways. Computation was guided by simultaneous progress toward three objectives: SGC grouping of identical functions (“spacing”), minimal mutation to reach the SGC (“distance”), and SGC’s minimal PR differences between codons related by single mutation (“dPR”). Thus, definitive origin information, the structure of the SGC, is combined with a coherent goal: a correct pathway for code emergence must yield the SGC. The major result is that an SGC-like coding table evolves easily, from small independent groups of codes (Fig. 9A, 10A), with no requirement for exotic events.

### The effective mechanism

The most rapid and accurate SGC evolution consigns translation initiation and termination to distinct, later evolutionary events; implements wobble after early non-wobbling code assignments; uses predominantly SGC-like, possibly stereochemical, assignment of sense codons and exploits coevolutionary mutational capture with assignments that conserve polar requirement.

### Wobble is inevitable in code descent

Wobble’s capture of third position mutation is required to emulate the SGC, but it is a double-edged sword. By extending initial triplet assignments to related wobbles, it decisively increases order. Such order is visible in initial spacing, distance and dPR progress arising from SGC-like triplets (Fig. 5B versus Fig. 5D, initial points, upper left). Nevertheless, subsequent evolution of a complete wobble code (22 encoded functions) is surprisingly prolonged (Fig. 5A); this is completion complexity. Continuous wobble’s slow evolution also allows destructive effects on pre-existing spacing and dPR order (Fig. 5C). Spacing progress is the exception, sustained at a moderate level by wobble’s persistent closely-spaced identical assignments (Fig. 5B).

### But non-wobble evolves to completion faster

Non-wobble code evolution contrasts strikingly with wobble: initial non-wobble allows quick code completion (Fig. 5C). However, because initial SGC triplet assignments are less effective, and wobble’s intrinsic enhancement of spacing (Fig. 5B) does not exist, spacing, distance and dPR still decline to near-random levels even during a non-wobbling code’s greatly shortened random-assignment era (Fig. 5D).

### Two wobble solutions

Non-wobble’s advantageous evolutionary rate and wobble’s ordering effects can combine if wobble was delayed, but immediately adopted into preexisting codes when translational advances, due to ribosomal evolution, made specific wobble possible. Moreover, because coevolution has a milder disruptive effect on spacing and distance order (Fig. 7A, 7C), and dPR can be enhanced by favoring conservation of polar requirement during biosynthetically related amino acid assignments (Fig. 7B), coevolution with intrinsic polar requirement matching (Coevo_PR) best balances the progress of a late-wobbling coding table (Fig. 7B). Twenty encoded functions are targeted to reduce completion complexity and because initiation and termination have distinct, unconserved mechanisms in life’s domains. Such late wobble yields coding that attains order close to SGC levels (Fig. 9A).

The second route to prompt wobble coding exploits a minority of wobble codes completed very early (continuous wobble; Fig. 6A). These reproduce SGC order well (Fig. 2C), and also exhibit similar sensitivity to random assignments (Fig. 8A, 8B). But while continuous wobble easily completes coding, it does not fill coding tables (Fig. 10, 11).

### Characteristics of code variation

To further resolve late wobble, the average 20-function late-wobbling coding table differs reproducibly from its exceptional subset in the SGC’s vicinity; with joint progress ≥ 0.9. Such differences objectively, quantitatively characterize favorable routes toward the SGC.

### Extended conclusion: SGC-like coding exploits simplicity

Selection of superior joint progress (Table II) dramatically increases resemblance to the SGC, as expected: from a mean of 0.7 – 0.8 to SGC-like levels of spacing, distance and dPR.

**Table II.** Average 20-function late wobble codes versus an excellent subset. Means for 10000 late-wobbling coding tables are shown down to the significant figure the same order as its sem. Evolution employed 10% random assignment, mutational capture of neighborhood triplets with coevo_PR, and late wobble at 20 encoded functions. There were 407/10000 tables with joint progress ≥ 0.9 (Fig. 7). Captions are defined in the supplementary lexicon.

| population | passages | decays | init | mut capt | wob init | spacing | distance | dPR |
| --- | --- | --- | --- | --- | --- | --- | --- | --- |
| Mean late wobble | 177 | 3.7 | 49.7 | 10.5 | 12.6 | 0.761 | 0.834 | 0.793 |
| joint progress $\geq 0.9$ | 115 | 1.8 | 42.3 | 6.3 | 15.1 | 0.98 | 0.96 | 1.02 |

Excellent coding appears in tables that have approximately half the average number of assignment decays. Superior coding also is achieved with about 60% the mutational captures occurring for an average code.

Initiations are also used more efficiently in coding tables which become more SGC-like. Only 85% of average initial assignments occur in the excellent code subset, and excellent codes reach 20 encoded functions in 65% of the average evolutionary duration.

That is: codes that resemble the SGC arise by chance simple routes, faster to complete. The favorable shortening of evolution in Table II is a smaller version of the ten thousand-fold superiority of 20-function late wobble compared to complete 22-function continuous wobble coding (Fig. 7A, 5A). Put another way - it seems unlikely that the SGC arose initially assigning codons an average of 300 times over, as implied for complete continuous wobble (Fig. 5A), or even 40 times/triplet on average, as for average 20-function continuous wobble (Fig. 5A). By comparison, 0.66 to 0.8 assignments/triplet to reach a near complete, late-wobbling code seems wholly credible (Table II).

### Routine unassigned triplets, late assignments, late wobble

Late unassigned triplets (Fig. 4A, 9, 10) support late assignments, deep into coding table evolution. This in turn is consistent with late-arising assignments of unique character for translation initiation and termination (Fig. 5A, 5C). Late unassigned triplets may also be advantageous if they provide for late advent of complex amino acids like tryptophan and methionine before encoding (Koonin and Novozhilov 2017).

Late unassigned triplets are even more pertinent for late-wobble advent. In fact, late wobbles are the exceptional more frequent evolutionary event among SGC-like coding tables (Table II). Because excellent SGC resemblance arises with fewer initiations of all kinds, more late wobble is used to fill in superior coding tables. In Table II, an average of 15.1 triplets are newly assigned after wobble is introduced to a superior 20-function code.

### Code sensitivity to random assignments

SGC-like order requires that randomly assigned codons be stringently limited in number (Fig. 2A). This imposes a limit, <= 15% random assignment if one requires accessible SGC regularity in either late wobbling (Fig. 8A) or continuous wobbling codes (Fig. 8B). Such a limit is essential because good spacing, distance and dPR occur together (Fig. 1A, Fig. 2C) in the SGC. It is therefore noteworthy that, all findings together, 1 of 24 late-wobbling coding tables, or 1 of 16 continuously-wobbling coding tables approach SGC order (joint progress >= 0.9 in Fig. 7B, 7D). Limited randomness is required for a combinatorial reason, considered next.

### Finding the SGC: the combinatorial abyss

Required simplicity hints at a much greater hindrance. Finding the ‘universal’ SGC demands exquisite discrimination. For slightly idealized coding tables like these, with 64 triplets and 20 encoded functions, there are 20^64^ = 1.8 x 10^83^ ways to assign triplets to functions if unassigned functions are allowed. Thus, there are astronomical numbers of possible non-wobbling genetic codes.

The situation is “improved” somewhat by wobble ordering; there are 32 two-codon wobble triplet groups, as assigned here, and 20^32^ = 4.3 x 10^41^ ways of assigning wobble groups to 20 functions, again with unassigned functions allowed. This is a minimum for wobbling genetic codes, because non-wobbling assignments are not counted, and will add to complexity.

The SGC (Standard Genetic Code) is an exceptionally ordered entity (Fig. 1A). Starting from unthinkably diverse sets like these, the SGC cannot plausibly be reached by starting at an arbitrary place, and/or taking an arbitrary path. Such an event has a small probability, because even arbitrary, pervasively ordered SGC-like tables (Fig. 1A) are a minute selection of total code configurations (Fig. 2C).

Alternatively, it is 1.43 x 10^17^ seconds since the Earth aggregated from the early solar disc (Patterson 1956). It is rational to ask: even if a quick-starting, planetary-scale selection exists to reject whole orders of possible codes/second, can the 24 to 66 order-of-magnitude disparity between a random code search and time available for searching be spanned, and an SGC found?

It therefore seems very improbable that the genetic code arose by exhaustive comparison of alternatives. Instead, the combinatorial abyss must have been virtually circumvented. That is also the finding here, on independent grounds (Fig. 7, 9, 10). The present 10% solution, mandating that 90% of initiations, more or less, correspond to SGC assignments, confines a coding table to a negotiable vicinity in code space near the SGC (see “**Sensitivity to random assignments**”, above). The result is wholly dramatic: evolution need not distinguish 10^83^ or 10^41^ options; instead, close SGC relatives appear in populations as small as hundreds of independent codes (Fig.7B, 7D, 9A, 10A).

But the abyss abides. Completion complications (Fig. 5A, 5C) are a portent, partly due to late evolutionary rates (Fig. 3), but also to the distance between almost complete and particular complete codes. Off-scale evolutions in in Fig. 6A and 6B are surely lost to the hiss of randomness. Code sensitivity to random substitution (Fig. 8A, 8B) is a hint of the combinatorial abyss.

### Independent evidence for non-random assignments

Coding tables must emphasize SGC-like assignments (**Finding the SGC**, just above). A large amount of independent evidence supports such specific triplet association with cognate amino acids.

### Experimental RNA binding sites

The most recent account of binding selection (Yarus 2017b) reviews data for 464 amino acid binding sites, all of independent molecular origin, selected from random sequence RNAs *in vitro* for specific binding of 8 amino acids of varied chemical classes. These include sites for disparate amino acid side chains: for example, as for polar Arg (Janas et al. 2010) and hydrophobic Ile (Lozupone et al. 2003). When the smallest RNA binding sites (perhaps more accessible in a primitive milieu) are specifically selected, the cognate triplet/amino acid association is observed in four of four cases (Yarus et al. 2009). Initially randomized nucleotide tracts in the same RNAs that are *not* required for amino acid binding are used as controls, and randomization and statistical tests show that triplet concentration in binding regions is specific and exceedingly non-random (Yarus 2017b). Statistical analysis requires assumptions, so it is notable that statistical tests are not essential to the crucial conclusion. Simplest sites and their triplets are so prevalent they are apparent when selected RNA sequences are simply aligned to reveal conserved sequences (for example, see L-Trp sites in: Majerfeld and Yarus 2005).

In total, comparisons of 7137 sequenced ribonucleotides within binding sites and 14,801 accompany control nucleotide sequences find that cognate triplets, whose nucleotides are essential to binding function, appear exceptionally often in amino acid binding sites. Thus selection reveals seven cognate anticodon triplets and two cognate codons *within* newly selected binding sites for six of eight tested amino acids. The two negative cases (L-Leu and L-Gln) are also among the least well explored; that is, those that yielded few sites for examination.

Further, a related tendency has been found in selected, specific RNA binding sites for peptide, like His-Phe (Turk-Macleod et al. 2012) when affinity for both side chains is demanded. When these experimental stereochemical interactions are added to chemical models, consistency with the genetic code has been shown to be improved (Buhrman et al. 2013). These selection experiments, especially when combined, strongly support stereochemical interactions as a basis of primordial coding.

### Natural RNAs

Natural examples of coding triplet/cognate amino acid also exist, such as the *Tetrahymena* active center (Yarus and Christian 1989), and the *Sulfobacillus* guanidinium riboswitch (Breaker et al. 2017; Yarus 2017b). Both bind arginine congeners to structures containing arginine codons.

### Bioinformatic analysis of biostructures

Moreover, there are data suggesting that relations between amino acids and their cognate coding triplets are yet more general. Coding triplets in present RNA biostructures appear significantly related to their amino acids. Within crystallographically defined ribosomes, shortened distances appear between protein amino acids and cognate rRNA triplets (Johnson and Wang 2010). Most remarkably, when mRNA sequences are examined across complete genomes (Polyansky et al. 2013), their cognate peptide sequences show significant correlations with mRNA sequences, consistent with amino-acid/RNA chemical interrelations. Such interrelations and their potential peptide/mRNA interactions persist even for accessible surfaces of folded proteins (Beier et al. 2014).

Thus: five independent arguments using data of varied types, point to stereochemical SGC assignments. Such assignments are required to order the SGC (Fig. 8A, 8B), they are required to find the SGC in the combinatorial abyss (**Coding history**, above), chemical interaction between RNA binding sites with essential triplets and their cognate amino acids has been selected, measured and characterized, parallel interactions are observed in natural RNAs, and bioinformatic analyses find a wide-ranging amino acid-codon relations consistent with such interactions.

### Unfamiliar mechanisms in coding history

Present models include events not usually discussed. Wobble captures a part of the underlying mutational pattern, and can strongly stimulate code order (Fig. 5B, 5D). Decays and reassignments are infrequently accorded key roles in coding history. But here, they are routine. Because their inclusion allows evolution of the SGC (Fig. 9, 10), and they are chemically plausible, they can have had a role in SGC history. Moreover, each has a potential code function: for example, reassignments allow recovery if non-specific initiations are inconsistent with SGC order (Fig. 8A, 8B). Of triplets assigned during average code history (Table II), 69% are initial assignments (perhaps 90 % of these stereochemical), 15% are mutational captures by an assigned triplet, and 16% are late appearing wobbles. Moreover, assignments decay, on average, once for 16 assigned triplets. The beautiful order of the SGC (Fig. 1A) does not rule out a heterogeneous origin.

### Mixed mechanisms in coding history

Successful present coding history emphasizes specific initial assignments; for example, stereochemical interactions. But it also calls on order from wobble, coevolution and paralogy. Such a mixed basis presently seems required, because different mechanisms emphasize order of distinct kinds. For example, wobble supports all progress, but particularly spacing and dPR (initial points, upper left: Fig. 5B, 5D). Each assignment capture mechanism makes a distinctive contribution to code order (Fig. 7), thus mixed contributions are needed to reach a broadly ordered SGC (Fig. 1A). In addition, a mixed history is independently plausible (e.g., Yarus 2017b) because coevolution and paralogy both require pre-existing assignments, implying pre-existing stereochemistry and/or minor randomness. The beautiful order of the SGC (Fig. 1A) does not rule out a heterogeneous origin.

### More definitive coding history

Late wobble with 85-90% SGC-like assignment, coevolution with polar requirement selection accurately locate the SGC vicinity (Fig. 9A, 10A), but can this accuracy be improved? Yes, likely.

Only a restricted inventory of effects has been considered. For example, homogeneous, minimal models using constant rates for assignment, decay and mutational capture were analyzed here. Plausible, more complex possibilities have not been examined. For examples, the possibilities that amino acids were encoded in subsets (Grosjean and Westhof 2016). Such segmentation would be very consistent with **Plausible primordial acceptors** above, and should be tested. Perhaps transitions and transversions should be distinguished. Perhaps encoding was partially by RNAs, and subsequently by nucleoproteins (Koonin and Novozhilov 2017), or perhaps the SGC is a community’s consensus (Vetsigian et al. 2006).

More generally: “…eventually one would reach a point where no new amino acid could be introduced without disrupting too many proteins. At this stage the code would be frozen” (Crick 1968). Given its universality, the SGC’s origin lies in deep time, defining Crick’s point. An accurate pathway that reproducibly attained the SGC (Fig. 1A), by linking to the Crick freezing point, would provide a credibly complete code history for discussion.

### Bayesian convergence

An objective criterion exists for such refinement. From Bayes’ Theorem, the more likely mechanism explains more aspects of the current SGC (Yarus et al. 2005b). Importantly, a hypothesis need not necessarily slowly become more plausible, given more evidence. Instead, it can multiply its probability if it explains independent aspects of the code. Such a “Bayesian convergence” can rapidly reinforce a correct explanation. Convergence remains useful for events unimaginably remote in time and scale, like the origin of the genetic code (Yarus et al. 2005b).

Convergence points to late wobble. This conclusion rests on two findings; in convergence, the argument’s force is determined by their combined weight.

### Late wobble fills coding tables

Late wobble quickly yields excellent coding tables, almost complete and almost full (Fig. 9). By comparison, continuous wobble also quickly creates excellent, almost complete coding tables (Fig. 10), but not almost full ones (**A highly significant difference…** above; Fig. 11). Late wobble around 20 pre-encoded functions is almost sufficient to the SGC - it has unassigned triplets that approximately meet a requirement for later initiation and termination. In contrast, continuous wobble requires a yet-unknown way to assign ≈ 20 triplets (Fig. 11).

### Late wobble allows better SGC access

Continuous wobble creates a subtle, but reproducible, advantage in code order (cf. Fig. 1A). Continuous wobble’s better joint progress appears in Fig. 2C’s distributions, in better prevalence of joint progress in Fig. 7D versus 7B, and around 10% randomness in Fig. 8B versus 8A. However, this advantage is lost because continuous wobble reaches the SGC through a minority of an evolving code population (Fig. 6A). Coding must be both ordered and complete (Fig. 1A), and over the usable range for random substitution (Fig. 8B, 8A), such continuously wobbling codes are 3 to 4-fold rarer. Accordingly, SGC access appears 3 to 4-fold more probable via late wobble.

### Distribution fitness exploits primordial fluctuation

SGC evolution exploits ‘distribution fitness’; that is, a rigorous requirement met by a heterogeneous group with excellent upper-tail members (Fig. 2C, 7). Undirected primordial variability is not a barrier, but instead is the crux of SGC emergence. This idea bears elaboration because it parallels previous findings.

There is an efficacious route to inherited gene expression, which requires only already-known RNA reactions (Yarus 2017a). Evolution of chemical inheritance is facilitated by a highly disperse population, from which selection readily picks extremely functional members. The pivotal diverse event is ‘starting bloc selection’, meaning selection of individuals initiating a reaction. Early starters have uniquely disperse product amounts – they are exceptionally suited to simultaneous selection of their product and its inheritance in a prebiotic, gene-free chemical system. In fact, a new inherited chemical capability can emerge after only one selection, possibly only a few days after partially activated primordial ribonucleotides accidently encounter each other (Yarus 2017a, 2018).

The starting bloc reappears during descent of the genetic code. In Table II, SGC-like codes are early-appearing, averaging 65% the duration of average code evolutions. Thus, if codes resembling the SGC were selected at an early time, SGC-like early starters would be prominent. Special capability is synonymous with early function, the typical linkage in starting bloc selection.

In the third example, prebiotic chemical systems must change to become biotic ones, so one may ask: how did prebiota advance without genes? One answer is ‘chance utility’, in which reactant variation permits persistent, unexpected evolutionary outcomes. For example, it is not only possible, but in a fluctuating milieu can be routine, that a desirable reactant is selected despite a 100-fold excess of a destructive competitor (Yarus 2016).

### Prebiotic history required selection of favorable fluctuations

Thus: chance utility, starting bloc selection and distribution fitness solve notable evolutionary problems because primitive systems offer fluctuation and distribution. In this way, unregulated primordial chemistry is intrinsically suited to evolutionary change: toward non-Darwinian chemical progress, toward primordial inheritance and later, toward Darwinian appearance of the genetic code. A connection between distributions and productive change suggests that prebiotic evolution itself may be a tractable branch of statistical mechanics. Prebiotic history presents puzzles of the size and complexity of planets; but even such puzzles can yield quantitative, probable solutions.

## Methods

### Computation

All calculations were performed on a Dell XPC laptop with an Intel Core i9 64-bit processor @ 2.9 GHz and 32 GB of RAM, running Microsoft Windows 10, v. 1709. Usually computer data were imported into Microsoft Excel 2016 32-bit as tab-delimited files for further analysis and conversion to graphics.

### Modeling

The probabilistic coding table model was developed and run in console mode of the Lazarus Integrated Development Environment v.1.8.4, with the Free Pascal Compiler v.3.0.4 supplying run-time modules. Pascal source code, Ctable18d1.pas, configured for late wobble but capable of all calculations presented with slight adjustments, and RandSens, a Microsoft Excel file that illustrates post-simulation calculation of sensitivity to random assignments, are available on request.

Because of the speed of integer operations, coding tables were represented as arrays of integers. One code array is followed to a specified evolutionary end: e.g., full coding or complete coding, using whatever evolutionary rules are being investigated. Such arrays were translated into ordinary coding tables (as in Fig. 1, 12) using an alphabetically related dictionary, after evolutionary calculations. Analysis of populations of finished arrays yields the probability of SGC-like results.

Runs with varied numbers of passages suggest that the ≈ 900-line program run as above requires about 4 μsec for one passage through a coding table and about 20 msec for one evolution (dependent on passage complexity).

### Flow during one passage through a nascent coding table

Fig. 12 sketches the flow of operations during one passage through an evolving coding table. Both continuous and late wobble are described, but alterations for the latter are in notes (as for 4. Below). Notations rand and rando are uniformly-distributed Mersenne Twister random numbers, 0 ≤ number ≤ 1, chosen anew for each passage. *P* terms are probabilities, as defined in the next section of Methods.

1. A triplet is chosen with random integers 1 to 4 for all 3 array indices.
2. Is the chosen triplet already assigned to a function, or currently free?
3. If unassigned, random number rand determines whether an assignment will occur, as shown.
4. Random number rando determines whether assignments are to a random choice of the 22 possible assignments, or is drawn from an assignment pre-existing in the SGC. In Fig. 12 as drawn, assignments are not unique, but Crick wobbles are encoded whenever possible. Late wobble: For late Crick wobble evolution, assignments are unique, and Crick wobble is added at step 8, wherever possible.
5. The random floating point number rand determines whether an assigned codon will decay or will capture a neighborhood triplet (related by single mutation) for a function related to its current assignment, as exemplified in Fig. 12.
6. The 9 neighborhood assignments for a current triplet are searched to see if any are free for mutational capture. If > 1 is free, choose randomly; if none are free, go to 8.
7. Random number rand determines whether capture of a randomly-chosen free triplet will occur. Related assignment mechanisms utilize a variety of means, as in Fig. 7A, 7C.
8. This operation represents a *while* loop that determines whether a desired coding property has been attained: e.g., is the coding table full?
9. Selected properties, depending on the goal of the calculation, for 10^2^ to 10^6^ successive coding tables, are calculated and written to disk.
10. A 3-dimensional code array is displayed as a conventional coding table (Fig. 1, 9, 10), with operations on both codons in a Crick wobble group, as in this example, indicated. Coding tables with specified properties can also be detected, viewed onscreen, and saved during calculations. If adjacent unassigned codons are not available for Crick wobble, unique assignments are allowed. Late wobble: if late instead of continuous wobble is utilized, unique assignments are made to completion (at Step 8, Fig. 12), then wobble is added wherever possible.

### Rate constants and probabilities

The kinetic method used here can be justified by showing that probabilities of reaction per passage are equivalent to use of normal rate constants.

### Initiations

The relation between *P*_*init*_ and the related first order rate constant, *k*_*init*_ in passages^-1^, can be calculated by equating kinetic and probability equations (probability for selection times probability for subsequent reaction) for the overall rate of initiations/passage:

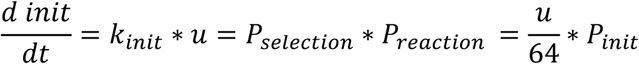

Where ***u*** is the number of ***u***nassigned triplets, and time is in passages. So

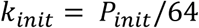

### Decays

A similar approach to a first order rate constant for assignment decay in passages^-1^ yields:

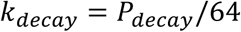

### Mutational captures

A second order rate constant, *k*_*mut*_, for mutational capture with units triplets^-1^ passages^-1^ must account for the probability that triplets neighboring an assigned triplet are so far unassigned, and can therefore be captured:

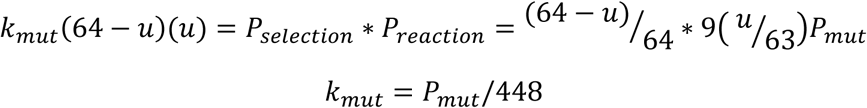

Where 9 (*u*/63) = *u*/7 is the expected number of unassigned triplets within the mutational neighborhood of a selected, assigned triplet.

### Controls

Controls suggested that randomly generated small mean probabilities derived from the Mersenne Twister algorithm in Free Pascal were accurate, that substitution of original experimental polar requirements for corrected ones would not materially change conclusions and that inclusion or exclusion of initiation/termination triplets from calculations where they are relevant would not substantially alter cited results. Transition probability variations alter coding, and have effects on order because they create the substrate for subsequent adoption into codes. But such alterations usually have smaller effects on order, and so have not been discussed in this first ms.

### Random spacing and distance values

The value for mean random mutational spacing and distance from an arbitrary triplet to other triplets in a random coding table can be calculated exactly from the fact that there are 9 triplets 1 mutation away from any initial triplet (its mutational neighborhood), 27 triplets 2 mutations away and 27 triplets 3 mutations away:

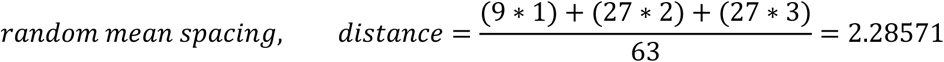

This is in excellent agreement with simulated values for 1000 randomized tables:

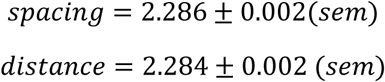

thereby validating programmed randomization and calculation of mean mutational distances.

## Supporting information

definitions for phrases with special meaning

## Acknowledgements

Many thanks are due Robin Dowell, John Heumann, Leslie Leinwand, Bill McClain and Jacob Stanley for helpful comments on early drafts of this work.

## Figure Legends

**Figure 9.** Sample coding tables from 600 late wobble, 20-function evolutions. The best observed, and sample codes having joint progress ≈1, ≈ 0.95 and ≈ 0.90 from late wobbling evolution are shown.

**Figure 10.** Sample coding tables from 600 continuous wobble evolutions. The best, and sample tables having joint progress ≈1, ≈ 0.95 and ≈ 0.90 at 200 passages are shown.

