## Supplementary material for "Evolution of the standard genetic code": definitions for phrases with special meaning

**Expressions used with a special meaning:**

**0 +/- 1 PR** – Paralogous mutational capture of a new codon, with the new assignment limited to amino acids separated by 0 or 1 places in the polar requirement series.

**0 +/- 2 PR** – Paralogous mutational capture of a new codon, with the new assignment limited to amino acids separated by 0, 1 or 2 places in the polar requirement series.

**0 +/- 3 PR** - Paralogous mutational capture of a new codon, with the new assignment limited to amino acids separated by 0, 1, 2 or 3 places in the polar requirement series.

**0 +/- 4 PR** - Paralogous mutational capture of a new codon, with the new assignment limited to amino acids separated by 0, 1, 2, 3 or 4 places in the polar requirement series.

**Bayesian convergence** – rapidly increasing probability for a hypothesis when multiple, particularly independent, observations are explained.

**Chance utility** – temporary or permanent change in a system without genetic inheritance; that is, a purely chemical system.

**Coevo** – coevolutionary mutational capture, with newly assigned amino acids having a biosynthetic relation to an existing one.

**Coevo_PR** – Coevo, but with new assignments constrained to favor a similar polar requirement.

**Completion complexity** – increased time and pathway complexity required for assignment of the last of 22 encoded functions.

**Continuous wobble** – wobble occurs continuously, throughout coding table history.

**Decay** – loss of a previously assigned triplet function.

**Distance** – progress value, varies from 0.0 to 1.0 as mean mutational distance between system triplets and identically assigned SGC triplets goes from random to the value for the SGC itself.

**Distribution fitness** - evolutionary success via varied individual fitness, the best of which are highly fit.

**dPR** – progress value varying from 0.0 to 1.0 as mean polar requirement difference between neighborhood triplets goes from random encoding to the (usually) smaller SGC value.

**Initiation** – assignment of 1 of 22 possible encoded functions to an unassigned triplet; abbreviated init.

**Late wobble** – evolutionary pathway in which assignments are to unique triplets, without wobble, then quick and complete adoption of a late-developing wobble mechanism.

**Mutational capture** – assignment of codon function related to that of an preexisting assigned neighborhood triplet, occurs only to unassigned triplets; abbreviated mut capt.

**Mutational neighborhood** – triplets related to a current one by single mutation, 9 neighbors per triplet.

**Paralogous, paralogy, paralogical** – mutational captures at random or by adaptation/selection, yielding amino acids with similar polar requirements assigned to related triplet codons.

**Passage** – one computational pass through a current coding table: passages are $\propto$ time.

**Pdecay** – probability that a chosen assigned triplet loses its assignment, per passage.

**Pinit** – probability of initial assignment for a random unassigned triplet, per passage.

**Pmut** – probability of mutational capture of an unassigned neighboring codon by an existing assignment, per passage.

**Prand** – probability that a new triplet assignment is random, rather than chosen from the SGC.

**Pwob** – probability of wobble if a codon triplet can be read either uniquely or via wobble, as for XYU.

**Spacing** – progress value varying from 0.0 to 1.0 as mean mutational distance between triplets with identical assignments goes from random to (usually) smaller SGC value.

**Starting bloc selection** – evolutionary success based on the fittest among a population being those just beginning the selected activity.
